# Delineating Structural Propensities of the 4E-BP2 Protein via Integrative Modelling and Clustering

**DOI:** 10.1101/2023.06.20.545775

**Authors:** Thomas E. Tsangaris, Spencer Smyth, Gregory-Neal W. Gomes, Zi Hao Liu, Moses Milchberg, Alaji Bah, Gregory A. Wasney, Julie D. Forman-Kay, Claudiu C. Gradinaru

## Abstract

The intrinsically disordered 4E-BP2 protein regulates mRNA cap-dependent translation through the interaction with the predominantly folded eukaryotic initiation factor 4E (eIF4E). Phosphorylation of 4E-BP2 dramatically reduces eIF4E binding, in part by stabilizing a binding-incompatible folded domain (REF). Here, we used a Rosetta-based sampling algorithm optimized for IDRs to generate initial ensembles for two phospho forms of 4E-BP2, non- and five-fold phosphorylated (NP and 5P, respectively), with the 5P folded domain flanked by N- and C-terminal IDRs (N-IDR and C-IDR, respectively). We then applied an integrative Bayesian approach to obtain NP and 5P conformational ensembles that agree with experimental data from nuclear magnetic resonance, small-angle X-ray scattering and single-molecule Förster resonance energy transfer (smFRET). For the NP state, inter-residue distance scaling and 2D maps revealed the role of charge segregation and pi interactions in driving contacts between distal regions of the chain (∼70 residues apart). The 5P ensemble shows prominent contacts of the N-IDR region with the two phosphosites in the folded domain, pT37 and pT46, and, to a lesser extent, delocalized interactions with the C-IDR region. Agglomerative hierarchical clustering led to partitioning of each of the two ensembles into four clusters, with different global dimensions and contact maps. This helped delineate an NP cluster that, based on our smFRET data, is compatible with the eIF4E-bound state. 5P clusters were differentiated by interactions of C-IDR with the folded domain and of the N-IDR with the two phosphosites in the folded domain. Our study provides both a better visualization of fundamental structural poses of 4E-BP2 and a set of falsifiable insights on intrachain interactions that bias folding and binding of this protein.

## 1. INTRODUCTION

Proteins are inherently dynamic and adopt conformations that range from very stable to completely disordered ^1^. An extreme case of protein polymorphism, intrinsically disordered proteins (IDPs) have been found to perform an increasingly diverse range of cellular functions, despite (or perhaps due to) lacking stable secondary and tertiary structure ^2^. Statistics of the human proteome revealed that nearly 60% of proteins contain stretches of greater than 30 residues of intrinsic disorder and ∼5% of proteins are completely disordered ^3^. IDPs are highly involved in cellular signalling and regulation, function as hubs of protein-protein interaction (PPI) networks ^4^, show unexpected mechanisms of PPIs ^5^, and are drivers of protein phase separation ^6^. They are particularly sensitive to post-translational modifications (PTMs), which can result in either stabilization or destabilization of transient secondary structures ^7^ and induce order-disorder ^8^ or disorder-to-order transitions ^9^. IDPs are enriched in many neurodegenerative and cancer pathways ^10^, but are challenging therapeutics targets due to the lack of stable binding pockets for small molecules ^11^.

Eukaryotic translation is a highly regulated process, with most mRNAs requiring interaction with the eukaryotic translation initiation factor (eIF4E) to be translated ^9, 12, 13^. The eIF4F complex is formed by assembly of eIF4E and eIF4G, which is subsequently recruited to the 40S subunit of the ribosome ^13^. The assembly of the eIF4F complex is inhibited by the intrinsically disordered 4E-BPs (eIF4E binding proteins), which compete with eIF4G for an overlapping surface of eIF4E ^14^.

The neuronal-specific 4E-BP isoform, 4E-BP2, modulates neuroplasticity, and impacts learning, memory formation ^15^, and autism spectrum disorders ^16^. 4E-BP2 binds eIF4E at both the canonical 54YDRKFLLDRR63 and a secondary 78IPGVT82 binding site; the canonical motif binds to eIF4E in a helical motif on the same convex surface as eIF4G ^14, 17^, while the secondary binding site is more dynamic and binds to the lateral surface of eIF4E ^18^.

Hierarchical phosphorylation of 4E-BP2 at residues T37, T46, T70, S65, and S83 results in the five-phosphorylated (5P) state and decreases the affinity of the 4E-BP2:eIF4E complex by ∼4000-fold compared to the non-phosphorylated (NP) state, via the formation of a 4-stranded β- sheet structure from residues 18-62 ^9, 19^. The initial two phosphorylations at residues T37 and T46 result in a ∼100-fold decrease in eIF4E affinity, while the additional phosphorylations in the C-terminal intrinsically disordered region (C-IDR) cause a further ∼40-fold decrease ^9, 19^.

Because of this, interactions with the C-IDR containing the additional three phosphosites were proposed to enhance stability of the folded β-sheet structure (which would reduce binding). In order to support this hypothesis or otherwise explain the enhanced stability/reduced 4E binding, structural models of full-length 4E-BP2 in both phosphostates are required.

The free energy landscapes of IDPs are typically shallow but not featureless, with local energy minima corresponding to transient secondary and tertiary structural biases which confer functional attributes ^20,21,22^. The potentially vast number of relevant structures makes the experimental and computational characterization of IDPs difficult. Modelling them necessitates a framework of sufficient complexity to capture relevant features, while avoiding being too large to be computationally intractable. IDPs are often modelled as conformational ensembles, which are a set of 3D structures (having x,y,z coordinates of each atom) with associated weights ^23^.

Data from nuclear magnetic resonance (NMR), small-angle X-ray scattering (SAXS), and single-molecule Förster resonance energy transfer (smFRET) can be used to refine a starting pool of conformations by imposing agreement with the experimental data ^24, 25^. Different experiments are sensitive to different length scales and timescales, with different degrees of time-averaging and ensemble-averaging. This is a heavily under-determined inverse problem, as the experimental restraints available are vastly insufficient to determine a unique conformational ensemble.

Several approaches have been applied to generate disordered conformational ensembles, such as Trajectory Directed Ensemble Sampling (TraDES) ^26^, flexible-meccano ^27^, IDPConformerGenerator ^28^and FastFloppyTail (FFT) ^29^. TraDES generates conformers by first building the backbone from Φ/Ψ angles sampled from a non-redundant set of structures from the PDB, geometric restraints and a Leonard-Jones type potential avoid steric clashes. Flexible-meccano samples amino acid specific Φ/Ψ potential wells from a compilation of non-secondary structure (loop) elements derived from the PDB. IDPConformerGenerator samples phi, psi and omega torsion angles from the PDB for various fragment lengths, and with different secondary structural biases, including based on experimental NMR chemical shifts. FFT is a PyRosetta based method that samples three-residue fragments from the PDB with a bias towards loop regions.

Optimization methods such as ENSEMBLE ^30^, Extended Experimental Inferential Structure Determination (X-EISD) ^31^, and Bayesian Maximum Entropy (BME) ^32^ reweight or select a subset of the initial conformational ensemble so that back-calculated biophysical observables match their experimental counterparts. The ENSEMBLE method uses pseudo-energy terms to quantify agreement between computation and experiment, where deviation from the initial ensemble is not being penalized. In contrast, X-EISD and BME methods use Bayesian frameworks that account for uncertainties in both experimental data and back-calculators. For example, BME treats the experimental data as time-/ensemble- averages and reweights the prior ensemble such that it agrees with experiments while maximizing the relative Shannon entropy. In this way, confidence is given to both the prior ensemble and the experimental data to prevent overfitting.

Arranging conformations into groups that share structural similarities, i.e., clusters, can lead to better visualization of heterogeneous IDP ensembles and help formulate structure- function relationships ^33^. The high degree of conformational disorder makes traditional similarity measures that require atomic superimposition of conformers ill-suited for IDPs ^34^. Conversely, a similarity criterion based on inter-residue alpha-carbon (Cα) Euclidean distance can be applied in agglomerative hierarchical clustering, which was shown to be a useful tool to characterize the heterogeneity of IDPs ^35^.

In this work, we applied the BME method ^32^ to optimize 4E-BP2 ensembles in both NP and 5P states that were generated by FFT ^29^. Agreement to experimental data such as the SAXS curve, two smFRET histograms, and C_α_/C_β_ Chemical Shifts (CS) for most of the chain (excluding residues within the folded domain in the 5P state), were imposed in the optimization procedure. An independent data set, the Paramagnetic Relaxation Enhancements (PREs) at several positions distributed along the 120-residue chain, was reserved for validation and for tuning the hyperparameters of the BME optimization.

Structural-based clustering suggests that NP 4E-BP2 predominantly samples four overall structural states. One of these clusters shares structural features with the eIF4E-bound state, indicating that some conformations contain preformed features than enhance the probability of complex formation upon collision with eIF4E. Contact maps of the 5P ensemble revealed pronounced interactions of the folded-domain phosphorylation sites pT37 and pT46 with N-IDR (residues 1-17), while contacts with the C-IDR were less frequent and more delocalized. 5P clustering analysis led to the separation of these interactions into four different clusters. This work describes highly probable structural poses and provides novel insights into the structure-function relation of a fascinating disordered protein that regulates translation initiation.

Importantly, it also provides specific ideas valuable for designing experiments to test the validity of these insights.

## 2. RESULTS

### Optimized 4E-BP2 ensembles

Motivated by the availability of structural data yet a lack of appropriate full-length computational ensembles of the 4E-BP2 protein, we calculated conformational ensembles consisting of 20,000 static conformers for both the NP and 5P variants. Our approach utilizes optimization and analysis methods that have been previously applied to model IDP ensembles ^29, 36^. A unique aspect of 4E-BP2 in comparison to other IDPs is the presence of a folded domain within the otherwise disordered 5P phosophoform. In this hyperphosphorylated state, a four-stranded beta-fold domain spanning residues 18-62 is stabilized. Modelling such a case motivated our choice of the FFT conformer generator ^29^, which allows the N- and C-IDRs to be sampled separately while maintaining folded domain poses derived from solution NMR experiments ^9^.

Optimization of the NP and 5P ensembles was performed with the BME method ^32^ using our previously published CS and smFRET data and new SAXS data (see Methods). [Note that, while sampling IDR tails and internal IDRs of proteins with folded domains is now possible within IDPConformerGenerator ^28^, it was not when our study began, nor was the current X- EISDv2 version with enhanced accessibility ^31^.] To validate and/or further optimize these ensembles, we evaluated their ability to reproduce experimental data that was withheld from the BME refinement process ^37^. As such, we further tuned the ensemble optimization using PRE data with its sensitivity to inter-residue contacts (< 25 Å).

For the NP ensemble (Fig. 1A), *χ^2^_Total_* decreases as the initial pool is reweighted and the effective fraction of conformations (*N*_eff_) decreases (see Methods 4.2). The decrease is initially steep, but then it levels-off with a markedly flatter slope below *N*_eff_ ≈ 0.6. The region of steep decrease is where the conformations that are least consistent with experimental data are essentially discarded, i.e., their weights go to zero. As the slope flattens, further optimization only marginally increases agreement with experiments and leads to overfitting. After an initial plateau, PRE RMSD follows a similar downward trend, although shifted to a lower *N*_eff_ range than *χ^2^_Total_*. To avoid overfitting, *N*_eff_ = 0.40> (*θ* = 35) was chosen at the “knee” point of the sampled PRE RMSD curve (see Methods 4.2) for the optimized NP ensemble.

**Figure 1.**
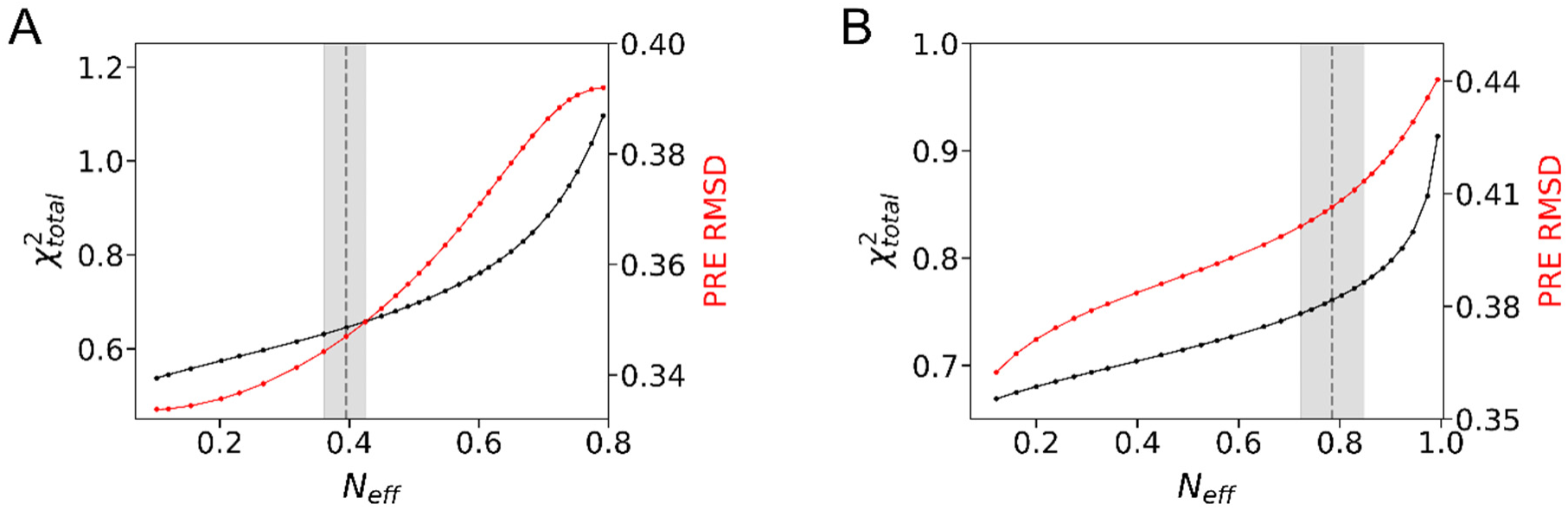
BME optimization for NP (**A**) and 5P 4E-BP2 (**B**) ensembles using FFT-generated prior pools with 20,000 conformers and imposing agreement with experimental data (SAXS, CS and FRET). A combination of fitting the restraints (*χ*^2^*_Total_*) and external validation (PRE RMSD, see SI 2.8) was used to determine the global fitting parameter *N*_eff_, indicated as dashed vertical lines and gray areas (see 4.2).

Similarly, for the 5P ensemble (**Fig. 1B**), increased conformer re-weighting leads to improved agreement with both the restraints incorporated within BME (decrease of *χ*^2^) and the external data (decrease of PRE RMSD). The knee points of the two curves are very close to each other, with the lower of the two, *N*_eff_ = 0.78 (*θ* = 27), being chosen for the optimized 5P ensemble. Fitting parameters of the BME-optimized ensembles are shown in **Table 1**.

**Table 1.** Fitness parameters and back-calculated global parameters for ensembles of NP and 5P 4E-BP2*

| | $N_{eff}$ | $\chi^2_{total}$ | $\chi^2_{FRET}$ | $\chi^2_{SAXS}$ | $\chi^2_{CS}$ | $R_g$ (Å) | $R_h^{HP}$ (Å) | $R_h^{KR}$ (Å) |
| --- | --- | --- | --- | --- | --- | --- | --- | --- |
| **NP**<br>**4E-BP2** | 0.40 | 0.64 | 1.03 | 0.67 | 0.62 | $28.7 \pm 0.1$ | $29.0 \pm 1.5$ | $23.5 \pm 0.1$ |
| **5P**<br>**4E-BP2** | 0.78 | 0.76 | 1.20 | 0.95 | 0.37 | $26.5 \pm 0.1$ | $26.8 \pm 1.5$ | $20.8 \pm 0.1$ |

\* Uncertainties of $R_g$ and $R_h$ are the weighted standard deviation of the mean of the ensemble distributions.

Optimization curves for each restraint are shown in Figs. S1-S2 in the SI and the initial and optimizing fitness parameters are displayed in Tables S1-S2. The effect of optimization can be visualized by the change in the distribution of conformer weights (Fig. S3). The NP distribution contains distinct outlier values that are well-separated from the bulk. In addition, 61% of the initial conformers have 95% of the weight in the optimized NP ensemble, while for 5P the fraction is much higher, 83%. This was perhaps expected since the 5P initial ensemble integrates atomic coordinates derived from the NMR solution structure of the folded domain (∼ 40 residues), and fewer residues required refinement.

**Table 1** also includes back-calculated global size parameters, radii of gyration and hydrodynamic radii (*R_g_* and *R_h_*), of the two optimized ensembles. The back-calculated *R_g_* values are close to those derived by Guinier analysis from the SAXS data (Fig. S11) and confirm that the 5P state is overall more compact than the NP state. The *R_h_* of the optimized NP 4E-BP2 ensemble, back-calculated using the Kirkwood-Riseman approximation (23.5 ± 0.1 Å), is closer to the value measured by FCS (24.8 ± 1.0 Å) ^20^ than the value back-calculated with HYDROPRO (29.0 ± 1.5 Å). Our results are consistent with a recent comparative study, where the Kirkwood-Riseman approach was shown to be a better predictor of experimental hydrodynamic radii of IDP ensembles and resulted in values ∼20% lower than HYDROPRO predictions ^38^. However, the Kirkwood-Riseman prediction for the 5P ensemble (20.8 ± 0.1 Å) is significantly smaller than the FCS-measured value (27.9 ± 1.1 Å) while the HYDROPRO prediction (26.8 ± 1.5 Å) is in better agreement. This discrepancy is perhaps not surprising, given that a significant fraction of the 5P protein (∼1/3 of the sequence) forms a stable fold, and HYDROPRO has been optimized to match the measured *R_h_* of folded proteins.

### Charge segregation and global compaction of NP 4E-BP2

Despite showing significant structural flexibility, IDPs have transiently sampled contacts due to intra-chain interactions such as hydrophobic^39^ ^40^, electrostatic ^41, 42^ and pi interactions ^43, 44^. Considering the global compaction of NP 4E-BP2 (see above), we asked whether there are indicators of non-local residue interactions in the optimized ensemble. As such, we analyzed the relation between mean inter-residue distances (*R_|i-j|_*) and residue separations (*|i - j|*), i.e., the Internal Scaling Profile (ISP). Distances were calculated as double averages, first for each conformer and then within the ensemble (Gomes JACS 2020). For comparison with a null-hypothesis lacking preferential interactions, we generated an ensemble consisting of 20,000 self-avoiding random coil (RC) conformations using TraDES ^26^ and computed its ISP curve.

Within the polymer physics framework, the ISP curve is typically fitted to the following power-law relation:

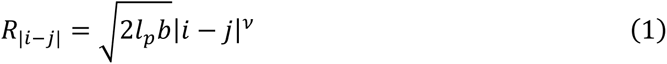

where *b* is the distance between bonded C_*α*_ atoms (3.8 Å), *v* is the Flory scaling exponent and the persistence length *l_p_* was fixed at *l_p_ =* 4 Å (see SI, Table S3 for fitting parameter values). This persistence length is commonly applied to model disordered proteins and has been shown to be applicable for unfolded and disordered proteins ^45^. The behavior of infinitely long homopolymer models representing the comparative strength of Protein-Protein Interactions (PPIs) vs Protein-Solvent Interactions (PSIs) converge for three distinct cases. A case in which PPIs dominate is termed the poor-solvent state (*v*∼0.33), PPIs being equal to PSIs is denoted as the *θ*-state (*v*∼0.5) and a chain with dominating PSIs is termed the good-solvent state, or the excluded-volume (EV) limit (*v*∼0.59).

To facilitate comparison, the ISPs of the optimized NP 4E-BP2 and TraDES RC ensembles are plotted together with the ISPs of the EV limit and the *θ*-state homopolymers (**Fig. 2A**). For sequence separations *10 ::; |i - j| ::; 40* the NP 4E-BP2 scaling resembles the TraDES RC ensemble (ν = 0.556), while for the largest separations, 100 ::; |*i - j*| ::; 120, the scaling exponent decreases only slightly (ν = 0.539). In the intermediate range, 60 ::; |*i - j*| ::; 95, the ISP curve flattens and undergoes a change in concavity, so it cannot be fit to a simple power-law dependence. In addition, intra-chain distances in the NP 4E-BP2 ensemble start to deviate from those in the TraDES RC ensemble for *|i- j| 2: 20* (**Fig. 2A**). Taken together, this suggests that scale invariance breaks down due to specific intra-chain contacts, which are also responsible for the high transient helical content spanning the entire chain ^14^ (Fig. S4).

**Figure 2.**
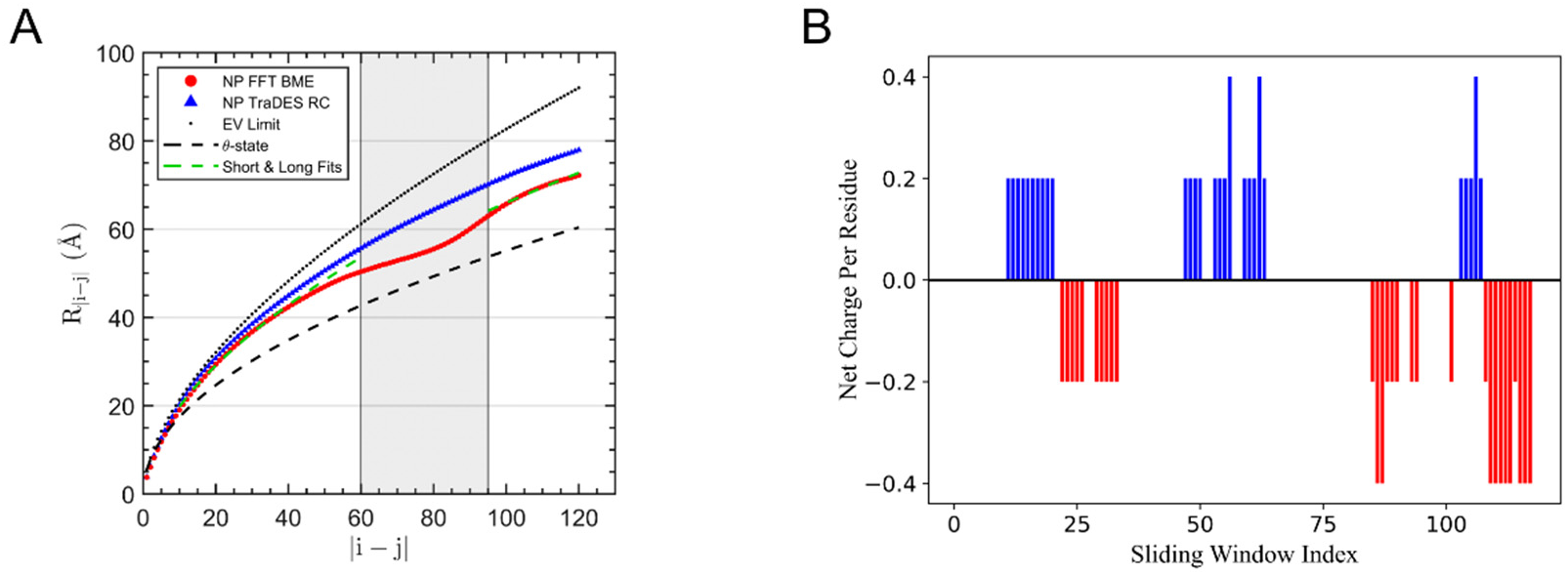
(A) Internal scaling profiles of the optimized NP 4E-BP2 ensemble (red), the TraDES random coil ensemble (blue), excluded-volume (black, dotted) and theta-solvent (black, dashed) homopolymers, and fits of the regions 10–40 and 100–121 to **Eq. 1** (green dashed). A concave region of the ISP curve, spanning residue separations of 60-95, is indicated by a grey shaded box. **(B)** Net-Charge-Per-Residue (NCPR) index calculated using a five-residue sliding-window; blue-positive, red-negative.

Charge segregation or patterning within a disordered chain can be quantified by the parameter *κ, 0 ::; κ ::; 1*, with the low limit corresponding to well-mixed charges and the high limit to positive and negative charges separated in the two halves of the chain ^46^, or by the sequence charge decoration (SCD) parameter ^47^. Das and Pappu tested the effects of charge segregation on the ISP behavior for a 50-residue model chain consisting of two oppositely charged residues that are distributed in patches of variable size across the sequence ^46^. They also observed a concavity “dip” in the ISP curves of model sequences, which became more pronounced with increasing *κ*. Interestingly, their model sequence with the closest *κ* value to NP 4E-BP2 (*κ = 0.1552*) has an ISP curve with a similar dip as our NP ensemble.

We evaluated various sequence-charge parameters using the Classification of Intrinsically Disordered Ensemble Relationships (CIDER) program ^48^ (Table S4). For example, the Net Charge Per Residue (NCPR) has been previously used to relate global dimensions of IDPs to electrostatic interactions^49^ ^42^. The NCPR map of NP 4E-BP2 (**Fig. 2B**) shows patches of oppositely charged residues in the sequence which may cause the dip in the ISP curve for *60 ::;|i - j| ::; 95* via electrostatic attraction. We identified three such attractive pairs: 11-24 (positive NCPR) with 85-98 (negative NCPR), 22-37 (negative NCPR) with 103-111 (positive NCPR), and 47-63 (positive NCPR) with 108-121 (negative NCPR).

To better visualize the proximity between different regions of the NP 4E-BP2 chain in our optimized ensemble, we constructed the 2D map of mean pairwise inter-residue Cα-Cα distance map normalized by each respective value from the RC ensemble (**Fig. 3A**). The most prominent region of compaction is centered between residues ∼20-40 and ∼80-100. The putative interacting regions based on NCPR analysis (**Fig. 3 B-D**) also contain hydrophobic, hydrogen-bonding and pi-containing residues. This suggests that transient contacts are formed through a combined effect of charge-based attraction with other physico-chemical interactions, potentially including the hydrophobic effect, hydrogen bonding and pi interactions.

**Figure 3.**
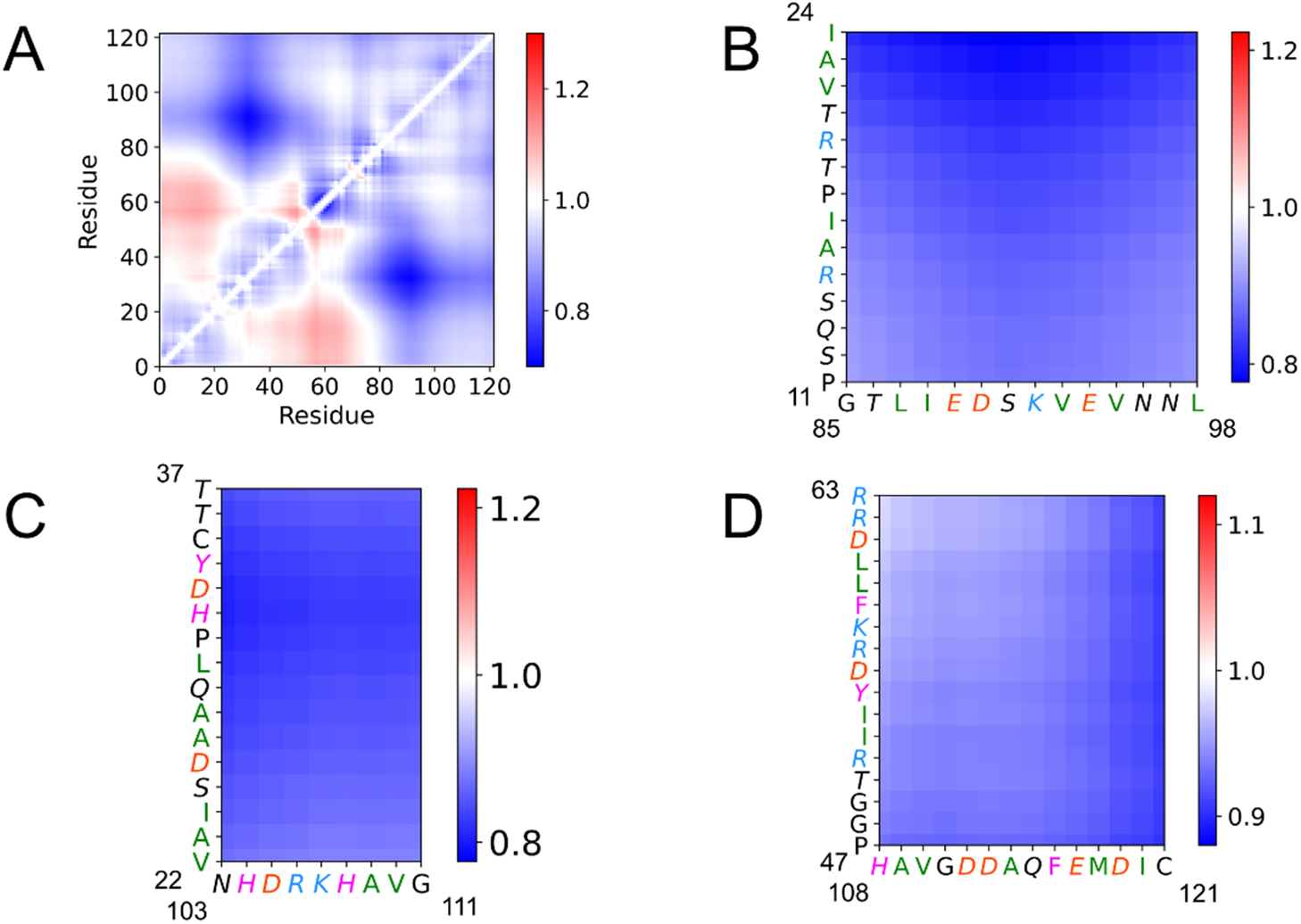
2D maps of mean inter-residue distances in NP 4E-BP2. (**A**) Distances in the BME-optimized ensemble normalized by the TraDES RC ensemble (red-expanded, blue-compacted). Zoom in the regions corresponding to pairs with opposite sign NCPRs (see **Fig. 2**): (**B**) residues 11-24 with residues 85-98, (**C**) 22-37 with 103-111, and (**D**) 47-63 with 108-121; residue color scheme: positive - blue, negative– red, hydrophobic – green, aromatic - magenta, hydrogen bonding – italic.

In particular, pi contacts between two tyrosines (Y34 and Y54) and two C-terminal lysines (K92, K107) and/or an arginine (R106) could contribute synergistically to the nonlocal interactions causing the dip in the ISP curve of NP 4E-BP2. Notably, for the first pair, the largest deviations from random coil expectations are located in residues of the positive NCPR selection and contain sites which are functionally relevant: the phosphoregulatory RAIP site (residues 15- 18) ^50^, and a region following the secondary binding site.

### Resolving non-local contacts that stabilize the folded domain of 5P 4E-BP2

Phosphorylation at residues T37, T46, S65, T70 and S83 induces the formation of a four-stranded beta-fold between residues 18-62 which sequesters the canonical eIF4E binding motif and is incompatible with binding ^9^. Phosphorylation is hierarchical. Initial phosphorylation at residues T37 and T46 leads to folding of a marginally stable domain, decreasing the eIF4E binding affinity by ca. 100-fold. Subsequent phosphorylation of the C-IDR at residues T70, S65 and S83 decrease the binding affinity by a further ca. 40-fold ^9^, primarily by stabilization of the folded domain and not by direct interactions with eIF4E. The non-cooperative folding/stabilization of this domain allows a graded inhibition of translation inhibition by phosphorylation induced tuning of the eIF4E:4E-BP2 affinity ^19^.

However, no structural models exist to provide detailed information on how the three additional C-IDR phosphorylation sites stabilize the folded domain, despite several experimental studies probing the properties of 5P 4E-BP2^9^ ^19, 20^. Molecular dynamics simulations have studied the formation of the four-stranded beta-fold but the N-IDR and C-IDR were omitted^51^ ^52^. NP 4E- BP2 contains significant transient α-helical structure, particularly between residues 49-67, partially pre-ordering the canonical helical eIF4E-binding element, and in the C-terminal region^14^. Phosphorylation at residues S37 and S46 switches this helical character to extended beta-like, and the additional C-IDR phosphorylations result in additional helical character in residues proximal to the canonical binding element as well as in the C-IDR, with pS65 having the largest effect ^19^. We examined our models to better understand stabilization of the fold by identifying potential C-IDR phosphorylation-induced stabilizing contacts between the folded domain and the rest of 4E-BP2 and potential destabilizing contacts present in the NP state that are abolished in the 5P state.

To evaluate 5P intra-chain interactions in the context of “topological” features imposed by the presence of a fixed folded domain, we compared the optimized 5P 4E-BP2 ensemble to the 5P coil ensemble (see SI 1.2). Similar to the NP analysis above, normalized pairwise inter-residue Cα-Cα distances reveal regions of compaction (*r_i,j_^norm^ < 1*) and expansion (*r_i,j_^norm^ > 1*).

Most inter-residue distances are closer in the 5P BME optimized ensemble compared to the 5P coil ensemble, with the closest contacts (besides those within the folded domain) involving residues of the folded domain with those of the N-IDR (**Fig. 4A**). Interestingly, the NP ensemble (**Fig. 3A**) showed greater distances between residues of the canonical binding motif^54^YXXXXLɸ^60^ and the N-terminus (residues 1-17), in contrast to the 5P state.

**Figure 4:**
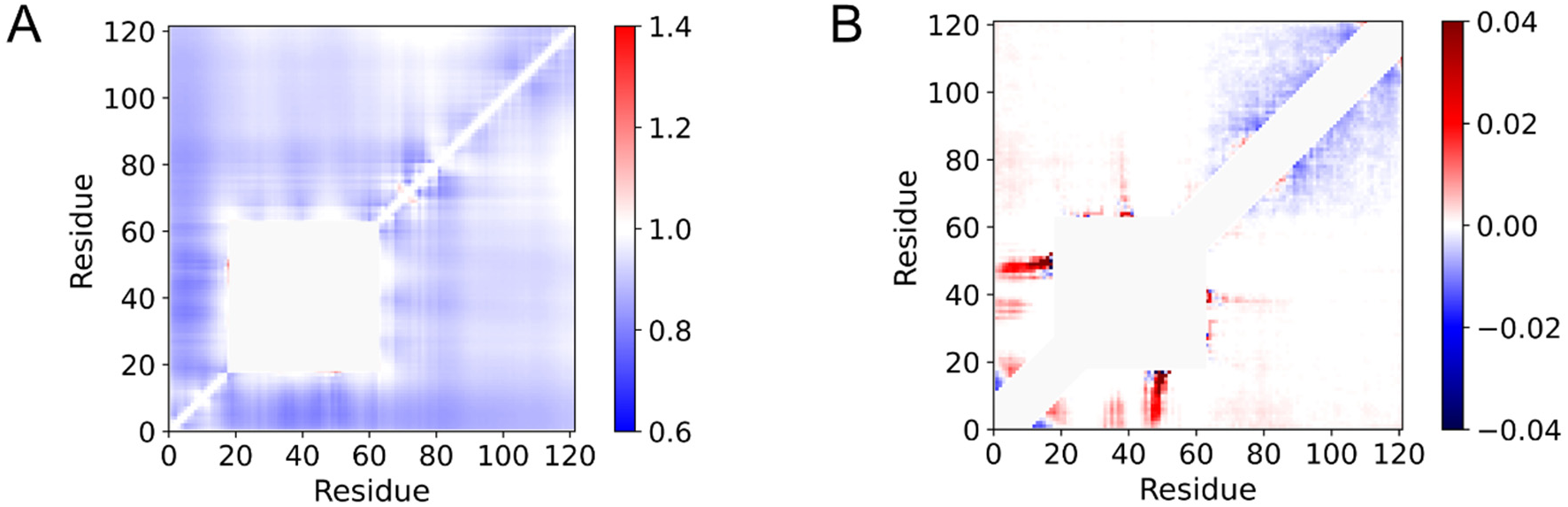
5P 4E-BP2 inter-residue distance and contact maps of optimized *vs.* coil ensembles. (**A**) 2D map of the mean inter-residue distances of the 5P 4E-BP2 optimized ensemble normalized by the 5P coil ensemble (red – expanded, blue – compacted). (**B**) Difference contact map obtained by subtracting the fractional degree of inter-residue contacts in the 5P coil ensemble from those in the BME-optimized 5P ensemble. Two residues are in contact if their C_α_ atoms are within 8 Å.

These changes are consistent with the observation that the chemical shift changes between the NP and the 5P state are the largest at the canonical binding site residues ^19^. In the NP state there are larger distances between the T46 phosphorylation site and all residues that will become the “N-IDR” upon phosphorylation than for the coil ensemble, and there are also larger distances between T37 and some residues in this N-IDR forming domain than in the coil (**Fig. 3A**). Conversely, in the 5P ensemble, the residues near phosphorylation sites pT37 and pT46 have distances that are the most reduced compared to the 5P coil ensemble. This can be seen more clearly by considering the difference contact map (**Fig. 4B**), where differences in fractional occupancy of inter-residue contacts between the optimized and the coil 5P ensembles are shown, with a contact defined as a Cα-Cα distance *< 8* Å (see SI 2.9). The areas of greatest positive contact difference are centered around the T37 and T46 phosphorylation sites and the N-IDR.

It has been proposed that C-IDR phosphorylation induces stabilizing contacts with the folded domain, possibly via electrostatic attractions between the C-IDR phosphate groups and the basic regions of the folded domain ^9, 19^. In our analysis, although the C-IDR is more compact than the random coil and shows sparse contacts with the folded domain, these contacts are not exclusive to the phosphorylation sites, implying that underlying interactions are of a mean-field nature. Instead, our results allude to a potential major role of the N-IDR in stabilizing the structure of the folded domain. The NCPR for 5P 4E-BP2 (see SI, Fig. S5) illustrates that the N-IDR is predominantly positive, while phosphorylation at T37 and T46 lead to a negative four charge difference in the folded domain.

A combination of electrostatic interactions between the basic N-IDR and the negative phospho-sites of the folded domain and between the basic parts of the folded domain and the negative phospho-sites in the C-IDR may increase the stability of the folded domain. At the same time, our analysis suggests that C-IDR phosphorylation disrupts the network of intramolecular interactions at regions far away from the phosphorylation sites with only small changes to the global dimensions, similar to other multi-phosphorylated proteins ^25, 53^ ^54^.

### Prominent 4E-BP2 structural states revealed by clustering

In contrast to stable folded proteins, IDPs feature a shallow and rugged free-energy landscape, without a pronounced global minimum. This facilitates fast conformational exchange, however weakly funneled landscapes exist for various IDPs^22^ ^55^ ^56^. Our previous NMR studies have shown that intra-chain interactions significantly affect conformational propensities of 4E- BP2 in different phosphorylation states^14^ ^9^ ^19^.

To better define non-local interactions impacting the 4E-BP2 structure, we applied agglomerative hierarchical clustering to partition the two optimized ensembles^57^ ^35^. The partitioning leads to a separation of global dimensions and shape, such as radius of gyration, end-to-end distance and asphericity (see SI 2.6, Figs. S6-S7). The dendrogram obtained from hierarchical clustering provides a visualization of the conformer amalgamation process (**Fig. 5A**). Motivated by the availability of experimental evidence for significant transient contacts, we sought to define states that are more likely to be populated in the function of this protein, since our computational models are optimized to agree with the experimental data.

**Figure 5.**
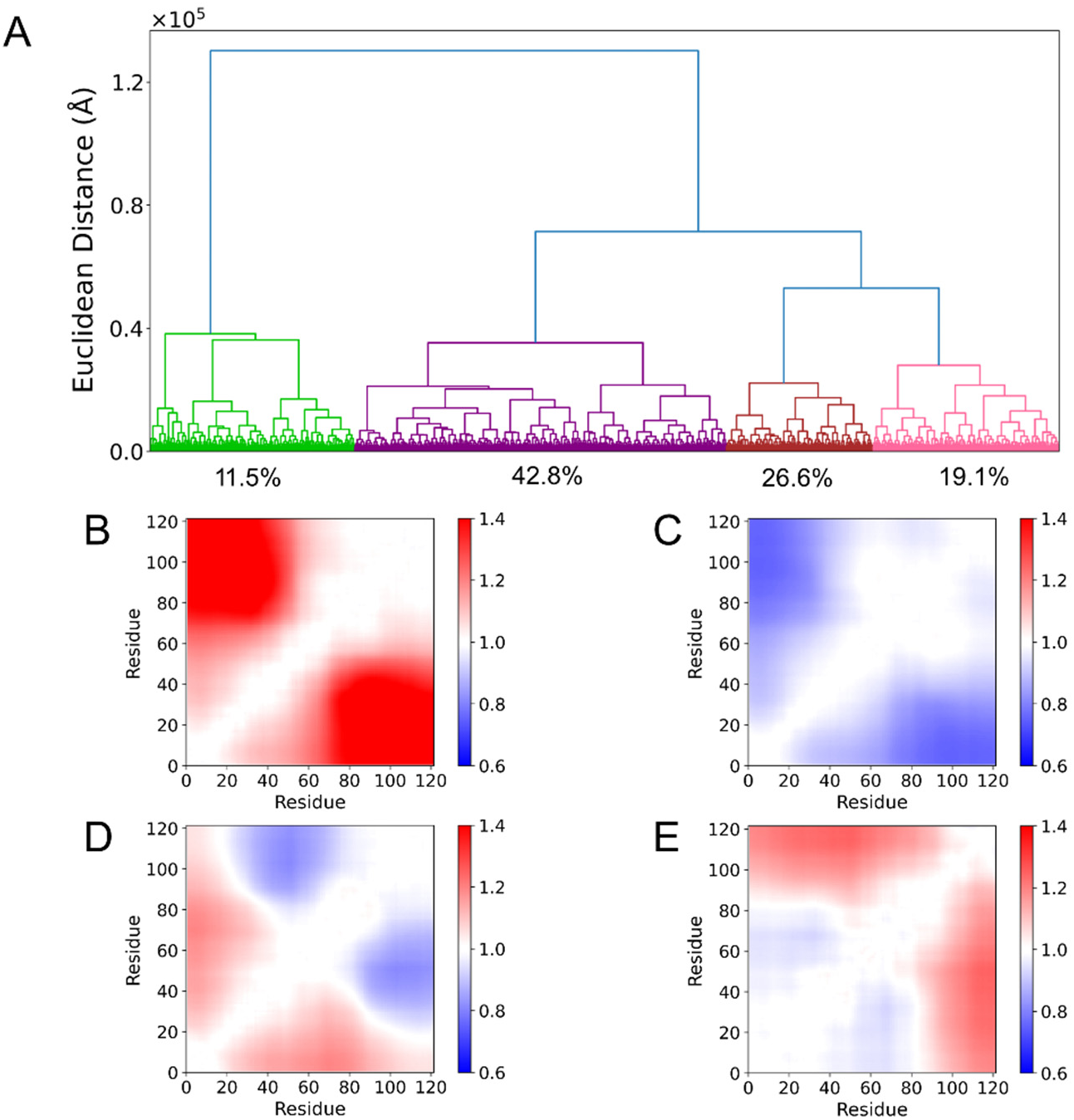
Agglomerative hierarchical clustering applied to the unrestrained NP 4E-BP2 ensemble. (**A**) Dendrogram showing the 4 resulting clusters: Cluster 1 (green), Cluster 2 (purple), Cluster 3 (brown) and Cluster 4 (pink). Inter-residue distance maps for each cluster normalized by the entire BME-optimized NP ensemble: (**B**) Cluster 1, (**C**) Cluster 2, (**D**) Cluster 3, and (**E**) Cluster 4.

The NP ensemble (unrestrained) partitions first into a small (23%, 4510 conformers) and a large (77%, 15490 conformers) cluster. The large cluster then splits twice before the cutoff criterion is satisfied (**Fig. S8A**), which brings the total number of clusters to four (**Fig. 5A**).

Upon reweighting the conformers with their BME-derived weights, the abundance of each cluster in the optimized ensemble is obtained (Table S5). Mean pairwise Cα inter-residue distances in each reweighted cluster were normalized by the corresponding distances for the optimized ensemble (**Figs. 5B-E**).

These maps confirm that the clusters have clearly distinct distributions of inter-residue distances, as expected since the dissimilarity metric used was a Euclidean distance between inter-residue distances in different conformers (see Methods 4.3). Note that such populations could not be trivially determined by analyzing the distribution of global parameters such as the radius of gyration (see SI, Fig. S9), underscoring the utility of clustering to disentangle coarse-grained structural propensities in a large and disordered protein ensemble.

Cluster 1 (green), whose fraction was reduced from 23% to ∼12% upon BME optimization, is the most expanded of all clusters (**Fig. 5B**). In particular, the N- and C-terminal regions are further apart, indicative of extended, quasi-linear poses. On the contrary, Cluster 2 (purple) is the most compact overall, while the other two clusters (3-brown, 4-magenta) have complementary distance maps, with a mixture of expansion and compaction compared to the full ensemble. Motivated by the growing literature on the binding mechanisms of IDPs^58^ ^59^ ^60^ and the expansion we previously captured between residues 32-91 and 73-121 of NP 4E-BP2 upon binding to eIF4E ^20^, we asked whether the expanded clusters were conformationally similar to bound-state structures.

To this end, back-calculated mean FRET values for each NP cluster were compared via a *z*-test (**Table 2**) to the experimental values obtained for the eIF4E-bound state, *E*_32-91_ = 0.26 ± 0.02 and *E*_73-121_ = 0.51 ± 0.02 ^20^. With this metric, Cluster 1 resembles the eIF4E-bound state, as it agrees within a 3*σ* tolerance level to the experimental values, in particular regarding *E*_32-91_. In contrast, the other three clusters have significantly higher *E*_32-91_ values than the bound-state, but instead agree within *3σ* tolerance with the apo-state value.

**Table 2.** Mean FRET efficiencies of NP clusters compared to apo- & bound-states values via a *z-*test*

| Cluster # | $E_{32-91}$ | $E_{73-121}$ | Bound<br>$z_{32-91}$ | Bound<br>$z_{73-121}$ | Apo<br>$z_{32-91}$ | Apo<br>$z_{73-121}$ |
| --- | --- | --- | --- | --- | --- | --- |
| Cluster 1 | 0.34 | 0.56 | 2.1 | 1.4 | 8.2 | 0.5 |
| Cluster 2 | 0.70 | 0.60 | 12.2 | 2.4 | 1.9 | 0.5 |
| Cluster 3 | 0.65 | 0.62 | 10.8 | 3.0 | 0.6 | 1.0 |
| Cluster 4 | 0.58 | 0.49 | 9.0 | 0.6 | 1.3 | 2.6 |

\* Uncertainties of back-calculated FRET efficiencies are $\pm 0.036$ (see Methods 4.4).

The presence of a sizeable cluster resembling the bound-state within the apo-state ensemble cannot be predicted *a priori*, as an ensemble could be split/clustered in many ways. Our results allude to a subclass of extended 4E-BP2 conformations that maximize attractive interactions with the eIF4E surface and initiate binding. At the same time, a large majority of conformers (88%) are not compatible with bound-state FRET. This suggests a hybrid model of binding, where both conformational selection and induced fit play a role, the latter being perhaps dominant for 4E-BP2. This remains an area of interest in the field, as both binding models have been proposed for disordered proteins ^61^ and combined binding mechanisms have also been described ^62^.

For the 5P ensemble, the normalized cutoff distance clearly levels off when increasing the number of clusters above *N* = 4 (see SI, Figure S8B). BME reweighting increases the population of the most expanded cluster from 20.1% to 30.6%, while reducing the population of the most compact cluster from 41.3% to 30.3% (Table S6). These observations agree with the overall expansion observed in 4E-BP2 upon hyper-phosphorylation using smFRET ^20^.

A more obvious clustering cutoff distance and a more balanced distribution of cluster fractions for the 5P ensemble compared to the NP ensemble suggest that the former energy landscape has fewer and deeper “structural wells” than the latter. The inter-residue distance maps reveal the complementarity of clusters (**Fig. 6B-E**). For instance, Cluster 1 mostly consists of conformers that are expanded throughout the entire chain, while the opposite is true for Cluster 2; similarly, Cluster 3 is compact in regions where Cluster 4 is expanded, and vice-versa.

**Figure 6.**
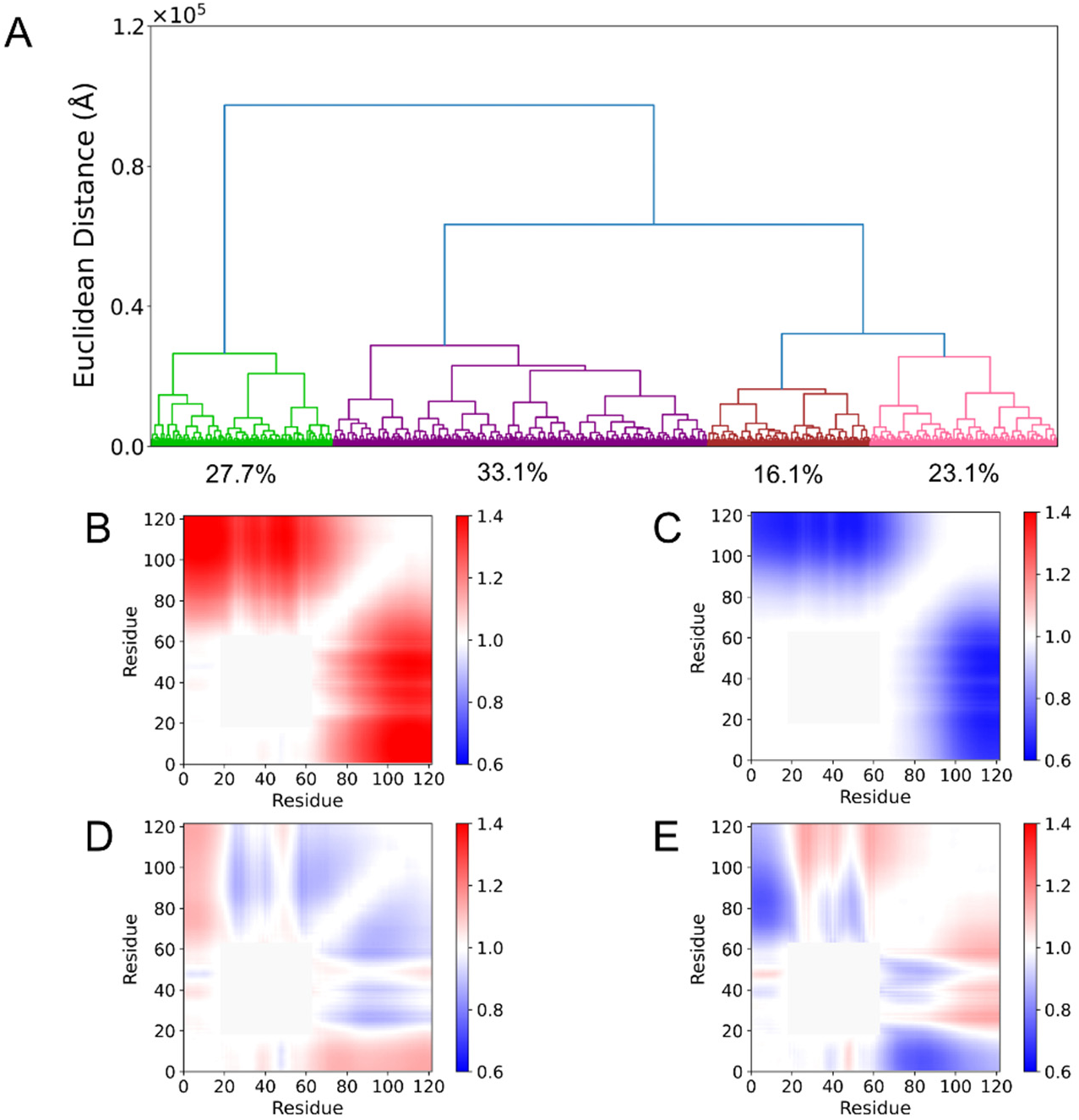
Agglomerative hierarchical clustering on the unrestrained 5P 4E-BP2 ensemble. (**A**) Dendrogram showing the 4 resulting clusters: Cluster 1 (green), Cluster 2 (purple), Cluster 3 (brown) and Cluster 4 (pink). Inter-residue distance maps for each cluster normalized by the entire BME-optimized 5P ensemble: (**B**) Cluster 1, (**C**) Cluster 2, (**D**) Cluster 3, and (**E**) Cluster 4.

Since interactions of disordered tails with the folded domain in 5P 4E-BP2 are thought to increase the stability of the fold ^19^, we then analyzed the abundance of intramolecular contacts at the cluster level (**Fig. 7**). Cluster 1 shows prominent contacts between the N-IDR and a segment around pT46 in the folded domain, while Cluster 2 shows a delocalized contact pattern between the C-IDR and the folded domain. Conversely, Clusters 3 and 4 contain prominent contacts between the N-IDR and pT46 and pT37 sites in the folded domain, respectively. We refer to Clusters 1 and 2 as C- Interaction Mode (CIM) clusters and Clusters 3 and 4 as N- Interaction Mode (NIM) clusters. To differentiate within the same mode, we denote Cluster 1 as CIM-off and Cluster 2 as CIM-on, while Clusters 3 and 4 are denoted as NIM-pT46 and NIM-pT37, respectively.

**Figure 7.**
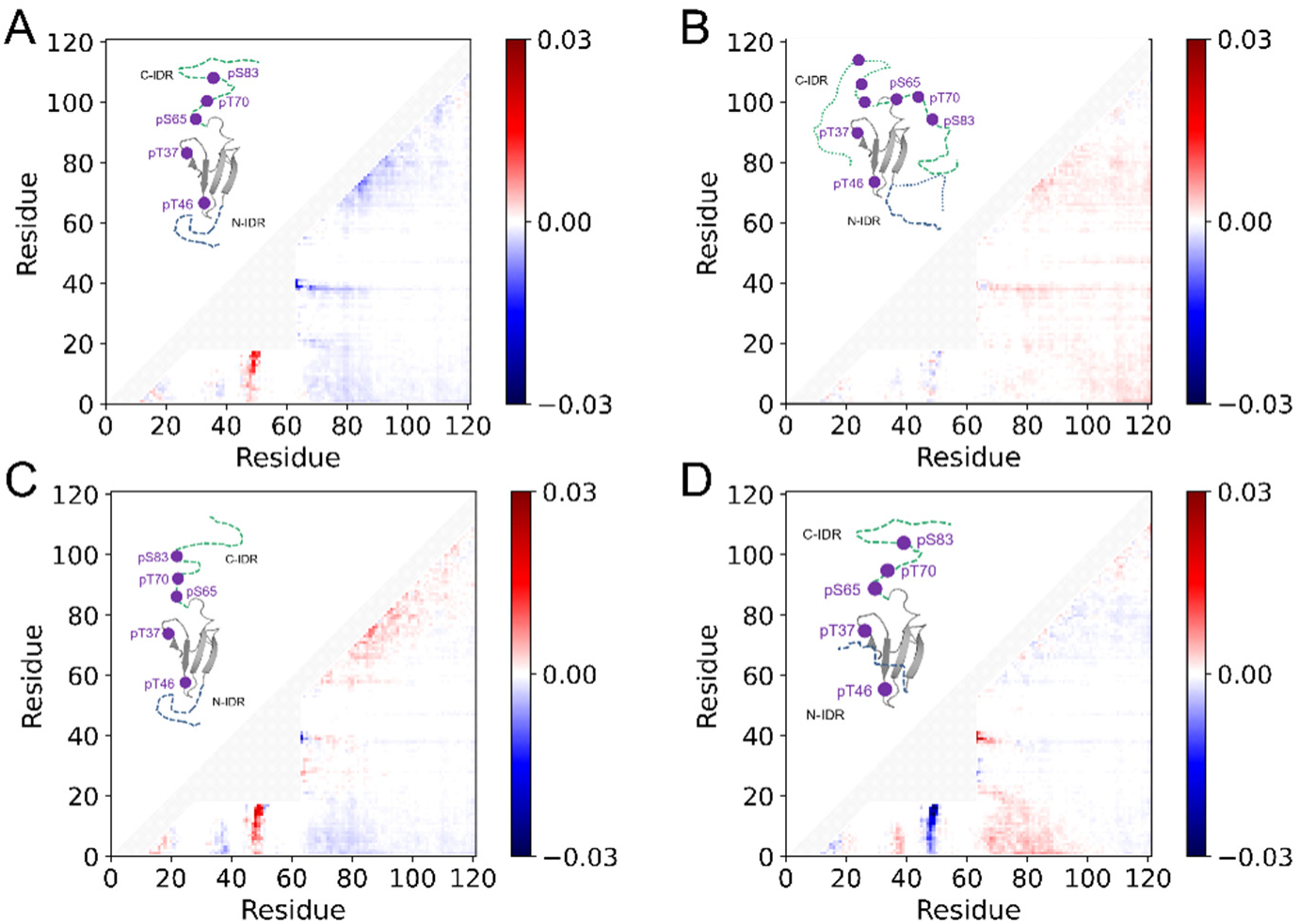
Difference contact maps obtained by subtracting the fractional level of inter-residue contacts in the entire BME-optimized 5P ensemble from those in each cluster. (**A**) Cluster 1 or CIM-off; (**B**) Cluster 2 or CIM-on; (**C**) Cluster 3 or NIM-pT46; (**D**) Cluster 4 or NIM-pT37. Representative conformations of each cluster are shown in the upper region of each panel. Two residues are in contact if their C_α_ atoms are within 8 Å.

These pairs of clusters represent two prominent modes of interaction within the BME- optimized 5P 4E-BP2 ensemble. The CIM-on cluster is enriched in contacts between residues 69-73 and 99-103 in the C-IDR with residues 14-19 and 10-13 in the N-IDR, respectively. Such contacts may be stabilized by attractive charge-based interactions as the aforementioned C-IDR residues have negative NCPR values and those within the N-IDR have positive NCPR values

(Fig. S5). To a lesser degree, contacts are formed between pT37 and pT46 of the folded domain and the entire C-IDR. This implies CIM-on conformations exhibit more contacts between the N-IDR and the C-IDR, which also brings the C-IDR in closer proximity to the folded domain for possible interactions. Such contacts are absent in the CIM-off cluster. Instead, CIM-off is enriched in contacts between a region near residue pT46 (47-51) and the entire N-IDR.

This N-IDR interaction with the folded domain (residues 44-53) is more prominent in the NIM-pT46 cluster, with the highest contact fractions between residues 6 and 48-49, 9-10 and 48, and 10-13 and 49. This cluster has minimal C-IDR contacts, resembling the CIM-off map. The NIM-pT37 cluster also has a high occurrence contact fraction between the N-IDR and the folded domain, except it is centered around residue pT37 and interacts with residues 1-11 of the N-IDR. The opposite sign NCPR values in these regions may aid in driving such contacts (Fig. S5). For these conformers, prominent contacts between the N-IDR and the C-IDR occur between residues 4-7 and 77-81, 4-6 and 66-69, and 1-3 and 102-105. This is similar to CIM-on conformers, suggesting that contacts occurring between the N- and C-IDRs facilitate interactions between the folded domain and the C-IDR.

## 3. DISCUSSION

Ensemble modelling of dynamic and/or disordered proteins is a growing area of research ^24, 25, 31, 63^, reflecting the increased awareness of their functional importance. Recently, we assessed the effects of various starting conformer pools and optimization methods for the integrative modelling of the disordered protein Sic1 ^37^. The quality of initial conformer pools had the highest impact in obtaining good agreement of the optimized ensemble with experimental data and is positively correlated with the *N*_eff_ value for the optimized ensemble. Using MD priors for the Sic1 protein, we found *N*_eff_ ≈ 0.75 ^37^, while a study that applied BME to MD simulations of the ACTR protein with CS restraints found *N*_eff_ ≈ 0.67 ^64^.

For 4E-BP2, the *N*_eff_ obtained for the optimized 5P ensemble is significantly higher than the value for the optimized NP ensemble, 0.78 *vs.* 0.40. This result is consistent with the aforementioned studies, as 5P conformers contain a significant folded fraction (residues 18-62). Although we included not one, but 20 different PDB structures to describe the folded domain ^9^, initial 5P conformers are much more restrained than initial NP conformers. For both phosphoforms, two smFRET efficiencies/distances were the most powerful restraints for the optimization procedure, while chemical shifts had the least impact, reflecting the high uncertainty from back-calculation of these values.

Interestingly, the NP ensemble appears to be more expanded overall than the 5P ensemble by *R_g_* (experimental, back-calculated) and *R_h_* (back-calculated) measures, while *R_h_* measured by FCS and two internal distances measured by smFRET show the opposite trend ^19, 20^. This may be a real effect reflecting different shapes/topologies of the two 4E-BP2 phosphoforms, or it may be an artefact due to limited sampling in the initial pools. Future studies will benefit from more accurate and diverse sampling of the conformational space, e.g., by better sampling at/around the five phosphorylation sites. In addition, more reliable back-calculators for NMR quantities (CS and PRE) and more smFRET distance restraints would significantly increase the confidence of the optimized ensembles.

Analysis of the optimized NP ensemble revealed a pronounced concavity in the inter-residue scaling profile for sequence separations on the order of 60-80 residues. This is likely caused by a combination of electrostatic charge mixing, hydrophobic interactions between residues 20-40 and 80-100, and pi interactions involving tyrosines Y34 and Y54 and C-terminal lysines and arginines.

Residues 19-28 show increased flexibility when bound to eIF4E ^14^. Our 2D distance maps point to interactions between these residues and residues in the C-terminal region controlling the non-local compaction of NP 4E-BP2. Binding to the surface of eIF4E may release these intra- molecular interactions, enhancing chain dynamics. This scenario would be consistent with our recent findings, in which we captured the expansion and increased local dynamics of 4E-BP2 upon binding to eIF4E ^20^. Furthermore, clustering analysis reveals only a minor sub-population (∼12%) that is “bound-state like”, also indicating major rearrangements of the chain when 4E- BP2 binds to eIF4E. smFRET experiments are currently under way probing different segments of the chain in the apo vs. the bound state. The new data will add important restraints to our modelling and help define the binding mechanism of this IDP system.

Difference distance maps reveal that the residues in the 5P folded domain show fewer contacts with the C-terminal region than they do in the NP state. Previously, we found evidence of a fast exchange between α-helical and β-strand conformations in the 2P state (pT37 and pT46), especially between residues 49 and 67 ^9^. Our results suggest that the addition of three phosphate groups in the C-terminal region may break the residue contacts that stabilize the α- helix, thus favoring the β-fold. Clustering analysis of the 5P ensemble captured four different modes of non-local interaction that define major topologies of the 5P state. This categorization of the ensemble’s heterogeneity hints at a mechanism for stabilization of the folded domain by the C-IDR in which the N-IDR acts both as a chaperone and an inhibitor.

This mechanism could act as follows: interaction of N-IDR with the folded domain driven by contacts with pT46 (present in CIM-off and more prominently in NIM-pT46) stabilizes a pose in which N- and C-IDRs are brought closer to each other. The average conformation then enters a NIM-pT37 average conformation where the pT46 contact is broken and the N-IDR forms contacts with the folded domain around pT37. This allows the C-IDR to form contacts with the starting residues of the N-IDR and permits the C-IDR to loosely interact with the starting residues of the folded domain. Finally, the N-IDR moves away from the folded domain as 5P 4E-BP2 enters the CIM-on state. Here, residues throughout the C-IDR form contacts around residues pT37 and pT46 after being led there by the N-IDR.

Additional ensemble models of the other 4E-BP2 phosphorylation states, in particular the two-fold phosphorylated state would contribute to unraveling this mechanism. Undoubtedly, combined efforts in improving the quality of the starting conformers, increasing the accuracy of back-calculators and obtaining new more restrictive experimental data will be instrumental in solving the fascinating molecular puzzle that is the 4E-BP2/eIF4E system.

## 4. METHODS

### 4.1 Conformer generation

4E-BP2 conformers were generated using the FastFloppyTail (FFT) algorithm, an optimized version of the Rosetta-based FloppyTail program which is ∼10 times faster and has enhanced accuracy via an improved fragment selection scheme ^29^ (see SI 1.1) . FFT has been applied to model inter-domain linkers ^36^ and several IDPs such as α-synuclein, Sic1 and the unfolded state of the drkN SH3 domain ^29^.

For NP 4E-BP2, we generated 20,000 conformers using FFT and a disorder prediction file created by the PsiPred DISOPRED3 web server ^65^. For the partially folded 5P 4E-BP2, we used FFT to sample the N- and C-termini (residues 1-17 and 63-121, respectively) with PsiPred DISOPRED3 disorder predictions. The folded domain (residues 18-62) consisted of the 20 lowest energy NMR-derived structures ^9^ and were fixed during FFT sampling of the IDRs. Each of the 20 PDB entries (PDB ID: 2MX4) were used with equal weight in generating the 5P 4E- BP2 ensemble (1000 structures per folded domain for a 20,000-conformer ensemble). The starting 5P structures had N- and C-terminal IDRs concatenated to the folded domain by using the “bond” function between pairs of carbon atoms in PyMOL ^66^. To create ideal bond lengths and angles while avoiding steric clashes, the “Idealize” and “Relax” Rosetta algorithms^67^ ^68^ were applied to the structures.

### 4.2 Ensemble refinement

We used the BME method ^32^ to refine the starting FFT conformational ensembles based on information supplied by experimental data. BME accounts for the uncertainty in estimating the confidence in the unrestrained FFT ensemble versus the experimental data by means of a tunable hyperparameter (*θ*). Given certain restraints, it holds that the most probable distribution compatible with the experimental data is the distribution of maximal entropy,^65^ ^69^. As such, conformer weights are tuned to minimize the following objective function:

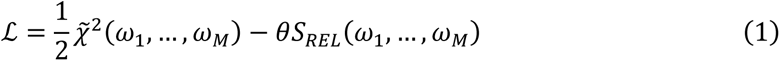

where *ω_i_* is the weight for conformer *i* in the reweighted ensemble, *M* is the number of conformers, *θ* is a hyperparameter which represents the degree of ensemble refinement and *χ̃*^2^ is the non-reduced chi-squared. Note that we refer to reduced chi-squared values (normalized by the number of degrees of freedom) without a tilde (*χ*^2^) and the non-reduced variant with a tilde (*χ̃*^2^). For more details, see SI section 2.1.

In the absence of a clear minimum on the optimization curve, L-curve analysis was applied to find the “knee” point using the *kneed* package in Python ^70^. The value of *N*_eff_ for both NP and 5P 4E-BP2 ensembles were determined by first removing concave down portions of the PRE RMSD curve plotted as a function of *N*_eff_ (see **Fig. 1**), and then sampling the spline interpolated plot at 1000 points uniformly throughout the curve. The “knee” point was then determined from the resulting discrete data to obtain the point of maximum curvature.

The points determined by the kneedle algorithm corresponds to the solid dashed lines in **Fig. 1** and the associated gray regions account for the variance across 5000-conformer replicate ensemble calculations of the same optimization (see SI, Tables S7, S8). More specifically, the gray region (on both sides of the dashed grey line) is the largest absolute difference between the point obtained via the above procedure for the 20,000-conformer ensemble analyzed in the main text and across the 5000-conformer replicates. This results in an *N*_eff_ uncertainty of *±*0.03 and *±*0.05 for the NP and 5P 4E-BP2 ensembles, respectively. The 5000-conformer replicate ensembles were generated by splitting the 20,000-conformer ensemble into four equally sized ensembles.

Upon refining ensembles with BME, the difficulty of fitting the mean FRET efficiency for NP 4E-BP2 labelled at residues 32 and 91 (*〈E〉_32-91_*) became apparent. Indeed, a large fraction of the prior ensemble must be discarded (*N*_eff_ = 0.03) to obtain good agreement with the experimental averages (*χ*^2^ *= 1.0*). The reason for this behavior is due to the BME protocol minimizing the sum: 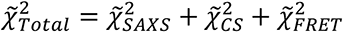, where each term in the sum is a non-reduced chi-squared. This means that experiments with many experimental datapoints, although not all independent (e.g., SAXS and CS), contribute much more to the total than smFRET. Hence, the optimization will be heavily biased towards reducing their *χ̃*^2^ values. To correct this, a hyperparameter controlling the weight of 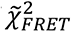 in the BME optimization was introduced *(Ω)*, modifying 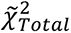 to the following form: 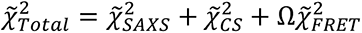, as implemented in our previous study ^37^. The hyperparameter *Ω* was tuned such that FRET was in good agreement with experimental values with negligible changes to the other restraints (see SI, Fig. S10).

Due to inaccuracies in both prior ensemble and experimental data, it is not clear what *θ* value should be selected for the most probable ensemble that fits all restraints. To resolve this issue, PRE data were not integrated as a restraint and were used instead to determine an optimal *N*_eff_ by choosing the “knee” point on the PRE RMSD curve that is uniformly sampled 1000 times for the full range of *N*_eff_ after spline interpolation. For comparison of experimental and back-calculated PRE NMR data, we have opted to compare ratios of intensities of peaks in the oxidized and reduced samples to back-calculated data using DEERPREdict ^71^; see below. We prefer comparing intensity ratios in contrast to a generally utilized strategy which converts PRE intensity ratios to distances ^72^. Such estimates are highly imprecise and, due to the required *r^-^*^6^ averaging, PRE distances act as a weak restraint where only a few conformers are needed to fit the data in order to achieve good agreement ^73^.

### 4.3 Hierarchical clustering

The NP and 5P 4E-BP2 ensembles were divided into sub-ensembles using hierarchical clustering using the Ward variance minimization algorithm ^74^. The distance metric for conformer (di)similarity is computed as the Euclidean distance in the 7260 – dimensional space where conformers are represented as matrices containing all non-degenerate pairwise inter-residue Cα- Cα distances. The distance *D_i,j_* between two conformers *i* and *j* is:

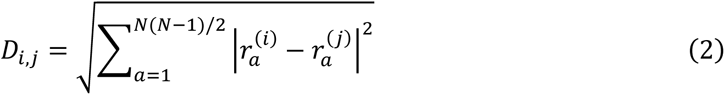

where *N* is the number of residues (121 in this case) and *r*(i) is the distance between Cα atoms of the *α^th^* residue pair for the *i^th^* conformer.

The dendrogram distance axis does not have a simple biophysical interpretation (see SI 2.4); we therefore transformed the dendrogram distance axis to a Euclidean distance between cluster means *(D_T_)* using the relation given by eq. S8 in SI, section 2.5 ^75^. We then divide this value by the square root of the number of non-degenerate inter-residue distance combinations to obtain an RMSD value of inter-residue Cα distances. We name this quantity, which is analogous to the atomic RMSD for protein structures (eq. S4), “normalized variance”.

To determine a cutoff for clustering, the number of clusters was plotted against the normalized variance (see Fig. S5), and L-curve analysis was applied to find the optimum number of clusters. This corresponds to 6 clusters for NP and 4 clusters for 5P. However, the three lowest populated clusters in the optimized NP ensemble (1.3%, 2.9%, and 7.3%) were combined into a single cluster to which these states are agglomerated.

### 4.4 FRET calculations

Back-calculated FRET efficiency, *〈E〉*, values of the IDP ensembles were computed via accessible volume simulations ^76 77^ using the *AvTraj* ^78^ and *MDTraj* ^79^ Python packages. We utilize dye parameters for Alexa488 and Alexa647 dye-linker systems documented previously ^80 81^. The back-calculated uncertainty was calculated by taking the difference between the mean FRET efficiencies in an ensemble computed using the lower and upper bounds of the Fӧrster radius, respectively ^25^. A back-calculated uncertainty of 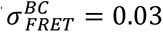 was computed for all ensembles. For use in BME, we added this uncertainty in quadrature with the experimental uncertainty 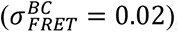, resulting in a combined uncertainty of 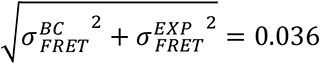 used for BME calculations.

### 4.5 SAXS data and calculations

The cloning, expression, purification and phosphorylation of 4E-BP2 was performed as previously described ^9, 14, 19^. A SAXSpace instrument with ASX autosampler (Anton-Paar GmbH, Austria) was used to conduct small-angle X-Ray scattering experiments. The SAXSpace was equipped with a long fine focus glass sealed copper tube using line collimation focus (40 kV/50 mA, Kα = 0.1542 nm), TCStage 150 sample holder and a 1D CMOS Mythen2 R 1K detector. 4E-BP2 protein samples at concentrations of 2-20 mg/mL were loaded into a 1 mm diameter quartz flow cell using the autosampler and six, 10-minute exposure frames were collected at 20°C under vacuum. Data was corrected for background scattering using sample buffer alone analyzed under the same conditions. SAXStreat software (Anton-Paar GmbH, Austria) was used to define the origin of the scattering curve, correct image distortion and convert the data to 1D scattering profiles. SAXSQuant (Anton-Paar GmbH, Austria) was then used to desmear the data.

The Pepsi-SAXS method ^82^ with default solvation parameters was used to back-calculate SAXS curves from IDP ensembles. Pepsi-SAXS is an efficient method which utilizes the multipole expansion scheme for scattering intensities and has been validated on more than 50 experimental SAXS scattering profiles. Only the experimental SAXS scattering intensity uncertainties were utilized in the BME optimization.

### 4.6 NMR data and calculations

The ShiftX program ^83^ was used to back-calculate secondary structure Chemical Shifts (CS) from the IDP ensembles. The ShiftX method can quickly compute backbone and sidechain ^1^H, ^13^C and ^15^N chemical shifts in for a single ∼100-residue conformer. All experimentally measured chemical shifts were employed in our BME calculations except in the 5P 4E-BP2 ensemble where phosphosites (65, 70, 83) and immediately subsequent residues (66, 71, 84) were excluded due to lack of functionality and inaccurate predictions indicated in the ShiftX output files, respectively. We also excluded 5P 4E-BP2 CS values assigned to residues within the folded domain (residues 19-61) since BME would not be able to refine the disordered conformer ensemble otherwise. Only back-calculation uncertainties of 0.98 and 1.10 were used for Cα and Cβ chemical shifts, respectively for BME calculations.

To generate PRE data for NP 4E-BP2, we first generated single-cysteine mutant constructs using a cysteineless version with C35 and C73 mutated to serines. Single cysteines were then introduced at positions 35, 65, 73, 91, 110 and 121 in order to attach a paramagnetic spin label at these positions. Proteins, that were labelled isotopically with 15N, were purified and a TEMPOL-maleimide (Toronto Research Chemicals) spin label was covalently linked as previously described ^19^. Two matched samples were made for each protein with the spin label in either an oxidized or a reduced state. Samples were oxidized or reduced by addition of either five-fold excess of TEMPOL (Toronto Research Chemicals) or 1 mM ascorbic acid, respectively. Prior to NMR experiments, the samples were buffer exchanged into a buffer containing 30 mM sodium phosphate, 100 mM sodium chloride, 1 mM EDTA, 1 mM benzamidine, pH 6 using argon purged buffers to maintain the oxidation state of the spin label.

For all samples, sensitivity-enhanced HSQC experiments ^84^ and unenhanced-NH-T2 experiments ^85^ were recorded at 20 °C on an 800 MHz Bruker spectrometer equipped with a triple-resonance cryoprobe. Relaxation delays for the T2 experiment were 7, 9, 14, 20, 26, 33, 41, 49, 59, 70, 82 and 95 ms, with the 14 and 59 ms points repeated for error estimation. A comparison of the T2 data and the ratios of the oxidized and reduced samples revealed highly similar trends. Though less rigorously quantitative, the peak intensities from the HSQC experiments were used as input for DEERPREdict (see below), because the T2 data and HSQC shared highly similar trends. PRE data for NP 4E-BP2 is included in the supplementary information, and the PRE data for 5P 4E-BP2 has been published previously ^19^.

The DEERPREdict program was used to back-calculate PRE intensity ratios ^71^. The parameters used were the same for both phosphoforms: total correlation time *τ_t_* = 0.5 ns, spin label effective correlation time *τ_c_* = 4 ns, total INEPT time *t_d_* = 10 ms, reduced transverse relaxation time *R_2_* = 6 Hz and proton Larmor frequency *ω_H_/2π* = 800.14. PRE data points for which both the spin label residue and residue to which it transfers were both in the folded domain were excluded in the analysis as they experienced little or no change. The metric chosen to quantify agreement is the root-mean-squared average over the root-mean-squared deviations between back-calculated and experimental PRE intensity ratios (PRE RMSD) (see SI 2.8).

## Supporting information

Supplemental infromation

## ACKNOWLEDGEMENTS

We thank John J. Ferrie from the Department of Chemistry at the University of Pennsylvania for advice on applying FastFloppyTail to generate 4E-BP2 conformers, as well as modifying the FastFloppyTail program so that it can include phosphorylated residues for the 5P state. This work has been supported by the Natural Sciences and Engineering Research Council of Canada (NSERC RGPIN-2023-04864 to C.C.G.) and the Canadian Institutes of Health Research (CIHR FND- 148375 to J.D.F.-K.).

## References

(1) Fisher, C. K.; Stultz, C. M. Protein Structure along the Order-Disorder Continuum. Journal of the American Chemical Society 2011, 133 (26), 10022–10025. DOI: 10.1021/ja203075p. Forman-Kay, J. D.; Mittag, T. From Sequence and Forces to Structure, Function, and Evolution of Intrinsically Disordered Proteins. Structure 2013, 21 (9), 1492–1499. DOI: 10.1016/j.str.2013.08.001.

(2) Berlow, R. B.; Dyson, H. J.; Wright, P. E. Functional advantages of dynamic protein disorder. Febs Lett 2015, 589 (19), 2433–2440. DOI: 10.1016/j.febslet.2015.06.003.

(3) Tsang, B.; Pritišanac, I.; Scherer, S. W.; Moses, A. M.; Forman-Kay, J. D. Phase Separation as a Missing Mechanism for Interpretation of Disease Mutations. Cell 2020, 183 (7), 1742-1756. DOI: https://doi.org/10.1016/j.cell.2020.11.050.

(4) Csizmok, V.; Follis, A. V.; Kriwacki, R. W.; Forman-Kay, J. D. Dynamic Protein Interaction Networks and New Structural Paradigms in Signaling. Chem Rev 2016, 116 (11), 6424-6462. DOI: 10.1021/acs.chemrev.5b00548. Dosztanyi, Z.; Chen, J.; Dunker, A. K.; Simon, I.; Tompa, P. Disorder and sequence repeats in hub proteins and their implications for network evolution. J Proteome Res 2006, 5 (11), 2985-2995. DOI: 10.1021/pr060171o.

(5) Borgia, A.; Borgia, M. B.; Bugge, K.; Kissling, V. M.; Heidarsson, P. O.; Fernandes, C. B.; Sottini, A.; Soranno, A.; Buholzer, K. J.; Nettels, D.;, et al. Extreme disorder in an ultrahigh-affinity protein complex. Nature 2018, 555 (7694), 61-+. DOI: 10.1038/nature25762.

(6) Martin, E. W.; Holehouse, A. S. Intrinsically disordered protein regions and phase separation: sequence determinants of assembly or lack thereof. Emerg Top Life Sci 2020, 4 (3), 307-329. DOI: 10.1042/Etls20190164. Pak, C. W.; Kosno, M.; Holehouse, A. S.; Padrick, S. B.; Mittal, A.; Ali, R.; Yunus, A. A.; Liu, D. R.; Pappu, R. V.; Rosen, M. K. Sequence Determinants of Intracellular Phase Separation by Complex Coacervation of a Disordered Protein. Mol Cell 2016, 63 (1), 72-85. DOI: 10.1016/j.molcel.2016.05.042.

(7) Baker, J. M. R.; Hudson, R. P.; Kanelis, V.; Choy, W. Y.; Thibodeau, P. H.; Thomas, P. J.; Forman-Kay, J. D. CFTR regulatory region interacts with NBD1 predominantly via multiple transient helices. Nat Struct Mol Biol 2007, 14 (8), 738-745. DOI: 10.1038/nsmb1278. He, Y. N.; Chen, Y. H.; Mooney, S. M.; Rajagopalan, K.; Bhargava, A.; Sacho, E.; Weninger, K.; Bryan, P. N.; Kulkarni, P.; Orban, J. Phosphorylation-induced Conformational Ensemble Switching in an Intrinsically Disordered Cancer/Testis Antigen. J Biol Chem 2015, 290 (41), 25090-25102. DOI: 10.1074/jbc.M115.658583.

(8) Banerjee, P. R.; Mitrea, D. M.; Kriwacki, R. W.; Deniz, A. A. Asymmetric Modulation of Protein Order– Disorder Transitions by Phosphorylation and Partner Binding. Angewandte Chemie International Edition 2016, 55 (5), 1675-1679. DOI: https://doi.org/10.1002/anie.201507728.

(9) Bah, A.; Vernon, R. M.; Siddiqui, Z.; Krzeminski, M.; Muhandiram, R.; Zhao, C.; Sonenberg, N.; Kay, L. E.; Forman-Kay, J. D. Folding of an intrinsically disordered protein by phosphorylation as a regulatory switch. Nature 2015, 519 (7541), 106-U240. DOI: 10.1038/nature13999.

(10) Uversky, V. N.; Oldfield, C. J.; Dunker, A. K. Intrinsically disordered proteins in human diseases: Introducing the D-2 concept. Annu Rev Biophys 2008, 37, 215-246. DOI: 10.1146/annurev.biophys.37.032807.125924.

(11) Biesaga, M.; Frigole-Vivas, M.; Salvatella, X. Intrinsically disordered proteins and biomolecular condensates as drug targets. Curr Opin Chem Biol 2021, 62, 90-100. DOI: 10.1016/j.cbpa.2021.02.009.

(12) Tait, S.; Dutta, K.; Cowburn, D.; Warwicker, J.; Doig, A. J.; McCarthy, J. E. G. Local control of a disorder-order transition in 4E-BP1 underpins regulation of translation via eIF4E. P Natl Acad Sci USA 2010, 107 (41), 17627-17632. DOI: 10.1073/pnas.1008242107. De Benedetti, A.; Harris, A. L. eIF4E expression in tumors: its possible role in progression of malignancies. Int J Biochem Cell B 1999, 31 (1), 59-72. DOI: Doi 10.1016/S1357-2725(98)00132-0.

(13) Sonenberg, N.; Hinnebusch, A. G. Regulation of Translation Initiation in Eukaryotes: Mechanisms and Biological Targets. Cell 2009, 136 (4), 731-745. DOI: 10.1016/j.cell.2009.01.042.

(14) Lukhele, S.; Bah, A.; Lin, H.; Sonenberg, N.; Forman-Kay, J. D. Interaction of the Eukaryotic Initiation Factor 4E with 4E-BP2 at a Dynamic Bipartite Interface. Structure 2013, 21 (12), 2186-2196. DOI: 10.1016/j.str.2013.08.030.

(15) Banko, J. L.; Merhav, M.; Stern, E.; Sonenberg, N.; Rosenblum, K.; Klann, E. Behavioral alterations in mice lacking the translation repressor 4E-BP2. Neurobiol Learn Mem 2007, 87 (2), 248-256. DOI: 10.1016/j.nlm.2006.08.012. Klann, E.; Sweatt, J. D. Altered protein synthesis is a trigger for long-term memory formation. Neurobiol Learn Mem 2008, 89 (3), 247-259. DOI: 10.1016/j.nlm.2007.08.009.

(16) Gkogkas, C. G.; Khoutorsky, A.; Ran, I.; Rampakakis, E.; Nevarko, T.; Weatherill, D. B.; Vasuta, C.; Yee, S.; Truitt, M.; Dallaire, P.;, et al. Autism-related deficits via dysregulated eIF4E-dependent translational control. Nature 2013, 493 (7432), 371-U113. DOI: 10.1038/nature11628.

(17) Peter, D.; Igreja, C.; Weber, R.; Wohlbold, L.; Weiler, C.; Ebertsch, L.; Weichenrieder, O.; Izaurralde, E. Molecular Architecture of 4E-BP Translational Inhibitors Bound to eIF4E. Mol Cell 2015, 57 (6), 1074- 1087. DOI: https://doi.org/10.1016/j.molcel.2015.01.017.

(18) Igreja, C.; Peter, D.; Weiler, C.; Izaurralde, E. 4E-BPs require non-canonical 4E-binding motifs and a lateral surface of eIF4E to repress translation. Nat Commun 2014, 5 (1), 4790. DOI: 10.1038/ncomms5790.

(19) Dawson, J. E.; Bah, A.; Zhang, Z. F.; Vernon, R. M.; Lin, H.; Chong, P. A.; Vanama, M.; Sonenberg, N.; Gradinaru, C. C.; Forman-Kay, J. D. Non-cooperative 4E-BP2 folding with exchange between eIF4E- binding and binding-incompatible states tunes cap-dependent translation inhibition. Nat Commun 2020, 11 (1). DOI: 10.1038/s41467-020-16783-8.

(20) Smyth, S.; Zhang, Z.; Bah, A.; Tsangaris, T. E.; Dawson, J.; Forman-Kay, J. D.; Gradinaru, C. C. Multisite phosphorylation and binding alter conformational dynamics of the 4E-BP2 protein. Biophys J 2022, 121 (16), 3049-3060. DOI: 10.1016/j.bpj.2022.07.015 From NLM Medline.

(21) Mittag, T.; Marsh, J.; Grishaev, A.; Orlicky, S.; Lin, H.; Sicheri, F.; Tyers, M.; Forman-Kay, J. D. Structure/Function Implications in a Dynamic Complex of the Intrinsically Disordered Sic1 with the Cdc4 Subunit of an SCF Ubiquitin Ligase. Structure 2010, 18 (4), 494-506. DOI: 10.1016/j.str.2010.01.020. Marsh, J. A.; Dancheck, B.; Ragusa, M. J.; Allaire, M.; Forman-Kay, J. D.; Peti, W. Structural Diversity in Free and Bound States of Intrinsically Disordered Protein Phosphatase 1 Regulators. Structure 2010, 18 (9), 1094-1103. DOI: 10.1016/j.str.2010.05.015.

(22) Papoian, G. A. Proteins with weakly funneled energy landscapes challenge the classical structure- function paradigm. P Natl Acad Sci USA 2008, 105 (38), 14237-14238. DOI: 10.1073/pnas.0807977105.

(23) Bonomi, M.; Heller, G. T.; Camilloni, C.; Vendruscolo, M. Principles of protein structural ensemble determination. Curr Opin Struc Biol 2017, 42, 106-116. DOI: 10.1016/j.sbi.2016.12.004. Jensen, M. R.; Zweckstetter, M.; Huang, J. R.; Backledge, M. Exploring Free-Energy Landscapes of Intrinsically Disordered Proteins at Atomic Resolution Using NMR Spectroscopy. Chem Rev 2014, 114 (13), 6632-6660. DOI: 10.1021/cr400688u. Marsh, J. A.; Forman-Kay, J. D. Ensemble modeling of protein disordered states: Experimental restraint contributions and validation. Proteins-Structure Function and Bioinformatics 2012, 80 (2), 556-572. DOI: 10.1002/prot.23220.

(24) Naudi-Fabra, S.; Tengo, M.; Jensen, M. R.; Blackledge, M.; Milles, S. Quantitative Description of Intrinsically Disordered Proteins Using Single-Molecule FRET, NMR, and SAXS. Journal of the American Chemical Society 2021, 143 (48), 20109-20121. DOI: 10.1021/jacs.1c06264.

(25) Gomes, G. N. W.; Krzeminski, M.; Namini, A.; Martin, E. W.; Mittag, T.; Head-Gordon, T.; Forman-Kay, J. D.; Gradinaru, C. C. Conformational Ensembles of an Intrinsically Disordered Protein Consistent with NMR, SAXS, and Single-Molecule FRET. Journal of the American Chemical Society 2020, 142 (37), 15697-15710. DOI: 10.1021/jacs.0c02088.

(26) Feldman, H. J.; Hogue, C. W. A fast method to sample real protein conformational space. Proteins 2000, 39 (2), 112-131. From NLM Medline. Feldman, H. J.; Hogue, C. W. V. Probabilistic sampling of protein conformations: New hope for brute force? Proteins-Structure Function and Bioinformatics 2002, 46 (1), 8-23.

(27) Ozenne, V.; Bauer, F.; Salmon, L.; Huang, J. R.; Jensen, M. R.; Segard, S.; Bernado, P.; Charavay, C.; Blackledge, M. Flexible-meccano: a tool for the generation of explicit ensemble descriptions of intrinsically disordered proteins and their associated experimental observables. Bioinformatics 2012, 28 (11), 1463-1470. DOI: 10.1093/bioinformatics/bts172.

(28) Teixeira, J. M. C.; Liu, Z. H.; Namini, A.; Li, J.; Vernon, R. M.; Krzeminski, M.; Shamandy, A. A.; Zhang, O.; Haghighatlari, M.; Yu, L.;, et al. IDPConformerGenerator: A Flexible Software Suite for Sampling the Conformational Space of Disordered Protein States. The Journal of Physical Chemistry A 2022, 126 (35), 5985-6003. DOI: 10.1021/acs.jpca.2c03726.

(29) Ferrie, J. J.; Petersson, E. J. A Unified De Novo Approach for Predicting the Structures of Ordered and Disordered Proteins. J Phys Chem B 2020, 124 (27), 5538-5548. DOI: 10.1021/acs.jpcb.0c02924 From NLM Medline.

(30) Krzeminski, M.; Marsh, J. A.; Neale, C.; Choy, W. Y.; Forman-Kay, J. D. Characterization of disordered proteins with ENSEMBLE. Bioinformatics 2013, 29 (3), 398-399. DOI: 10.1093/bioinformatics/bts701 From NLM Medline.

(31) Lincoff, J.; Haghighatlari, M.; Krzeminski, M.; Teixeira, J. M. C.; Gomes, G. N. W.; Gradinaru, C. C.; Forman-Kay, J. D.; Head-Gordon, T. Extended experimental inferential structure determination method in determining the structural ensembles of disordered protein states. Commun Chem 2020, 3 (1). DOI: ARTN 74 10.1038/s42004-020-0323-0.

(32) Bottaro, S.; Bengtsen, T.; Lindorff-Larsen, K. Integrating Molecular Simulation and Experimental Data: A Bayesian/Maximum Entropy Reweighting Approach. Methods Mol Biol 2020, 2112, 219-240. DOI: 10.1007/978-1-0716-0270-6_15 From NLM Medline.

(33) Appadurai, R.; Koneru, J. K.; Bonomi, M.; Robustelli, P.; Srivastava, A. Demultiplexing the heterogeneous conformational ensembles of intrinsically disordered proteins into structurally similar clusters. bioRxiv 2022, 2022.2011.2011.516231. DOI: 10.1101/2022.11.11.516231.

(34) Lazar, T.; Guharoy, M.; Vranken, W.; Rauscher, S.; Wodak, S. J.; Tompa, P. Distance-Based Metrics for Comparing Conformational Ensembles of Intrinsically Disordered Proteins. Biophys J 2020, 118 (12), 2952-2965. DOI: 10.1016/j.bpj.2020.05.015 From NLM Medline.

(35) Baul, U.; Chakraborty, D.; Mugnai, M. L.; Straub, J. E.; Thirumalai, D. Sequence Effects on Size, Shape, and Structural Heterogeneity in Intrinsically Disordered Proteins. J Phys Chem B 2019, 123 (16), 3462-3474. DOI: 10.1021/acs.jpcb.9b02575 From NLM Medline.

(36) Graham, T. G. W.; Ferrie, J. J.; Dailey, G. M.; Tjian, R.; Darzacq, X. Detecting molecular interactions in live-cell single-molecule imaging with proximity-assisted photoactivation (PAPA). Elife 2022, 11. DOI: 10.7554/eLife.76870 From NLM Medline.

(37) Gomes, G. W.; Namini, A.; Gradinaru, C. C. Integrative Conformational Ensembles of Sic1 Using Different Initial Pools and Optimization Methods. Front Mol Biosci 2022, 9, 910956. DOI: 10.3389/fmolb.2022.910956 From NLM PubMed-not-MEDLINE.

(38) Pesce, F.; Newcombe, E. A.; Seiffert, P.; Tranchant, E. E.; Olsen, J. G.; Grace, C. R.; Kragelund, B. B.; Lindorff-Larsen, K. Assessment of models for calculating the hydrodynamic radius of intrinsically disordered proteins. Biophys J 2023, 122 (2), 310-321. DOI: 10.1016/j.bpj.2022.12.013 From NLM Medline.

(39) Felitsky, D. J.; Lietzow, M. A.; Dyson, H. J.; Wright, P. E. Modeling transient collapsed states of an unfolded protein to provide insights into early folding events. Proc Natl Acad Sci U S A 2008, 105 (17), 6278-6283. DOI: 10.1073/pnas.0710641105 From NLM Medline.

(40) Yamada, J.; Phillips, J. L.; Patel, S.; Goldfien, G.; Calestagne-Morelli, A.; Huang, H.; Reza, R.; Acheson, J.; Krishnan, V. V.; Newsam, S.;, et al. A bimodal distribution of two distinct categories of intrinsically disordered structures with separate functions in FG nucleoporins. Mol Cell Proteomics 2010, 9 (10), 2205-2224. DOI: 10.1074/mcp.M000035-MCP201 From NLM Medline.

(41) Sizemore, S. M.; Cope, S. M.; Roy, A.; Ghirlanda, G.; Vaiana, S. M. Slow Internal Dynamics and Charge Expansion in the Disordered Protein CGRP: A Comparison with Amylin. Biophys J 2015, 109 (5), 1038-1048. DOI: 10.1016/j.bpj.2015.07.023 From NLM Medline.

(42) Bianchi, G.; Longhi, S.; Grandori, R.; Brocca, S. Relevance of Electrostatic Charges in Compactness, Aggregation, and Phase Separation of Intrinsically Disordered Proteins. Int J Mol Sci 2020, 21 (17). DOI: 10.3390/ijms21176208 From NLM Medline.

(43) Jo, Y.; Jang, J.; Song, D.; Park, H.; Jung, Y. Determinants for intrinsically disordered protein recruitment into phase-separated protein condensates. Chem Sci 2022, 13 (2), 522-530. DOI: 10.1039/d1sc05672g From NLM PubMed-not-MEDLINE.

(44) Vernon, R. M.; Chong, P. A.; Tsang, B.; Kim, T. H.; Bah, A.; Farber, P.; Lin, H.; Forman-Kay, J. D. Pi-Pi contacts are an overlooked protein feature relevant to phase separation. Elife 2018, 7. DOI: 10.7554/eLife.31486 From NLM Medline.

(45) Hofmann, H.; Soranno, A.; Borgia, A.; Gast, K.; Nettels, D.; Schuler, B. Polymer scaling laws of unfolded and intrinsically disordered proteins quantified with single-molecule spectroscopy. Proceedings of the National Academy of Sciences 2012, 109 (40), 16155-16160. DOI: doi:10.1073/pnas.1207719109.

(46) Das, R. K.; Pappu, R. V. Conformations of intrinsically disordered proteins are influenced by linear sequence distributions of oppositely charged residues. Proc Natl Acad Sci U S A 2013, 110 (33), 13392-13397. DOI: 10.1073/pnas.1304749110 From NLM Medline.

(47) Sawle, L.; Ghosh, K. A theoretical method to compute sequence dependent configurational properties in charged polymers and proteins. The Journal of Chemical Physics 2015, 143 (8). DOI: 10.1063/1.4929391 (acccessed 5/18/2023).

(48) Holehouse, A. S.; Das, R. K.; Ahad, J. N.; Richardson, M. O.; Pappu, R. V. CIDER: Resources to Analyze Sequence-Ensemble Relationships of Intrinsically Disordered Proteins. Biophys J 2017, 112 (1), 16-21. DOI: 10.1016/j.bpj.2016.11.3200 From NLM Medline.

(49) Mao, A. H.; Crick, S. L.; Vitalis, A.; Chicoine, C. L.; Pappu, R. V. Net charge per residue modulates conformational ensembles of intrinsically disordered proteins. Proc Natl Acad Sci U S A 2010, 107 (18), 8183-8188. DOI: 10.1073/pnas.0911107107 From NLM Medline.

(50) Wang, X.; Beugnet, A.; Murakami, M.; Yamanaka, S.; Proud, C. G. Distinct Signaling Events Downstream of mTOR Cooperate To Mediate the Effects of Amino Acids and Insulin on Initiation Factor 4E-Binding Proteins. Molecular and Cellular Biology 2005, 25 (7), 2558-2572. DOI: 10.1128/MCB.25.7.2558-2572.2005.

(51) Bomblies, R.; Luitz, M. P.; Zacharias, M. Molecular Dynamics Analysis of 4E-BP2 Protein Fold Stabilization Induced by Phosphorylation. Journal of Physical Chemistry B 2017, 121 (15), 3387-3393. DOI: 10.1021/acs.jpcb.6b08597.

(52) Zeng, J.; Jiang, F.; Wu, Y. D. Mechanism of Phosphorylation-Induced Folding of 4E-BP2 Revealed by Molecular Dynamics Simulations. J Chem Theory Comput 2017, 13 (1), 320-328. DOI: 10.1021/acs.jctc.6b00848.

(53) Gibbs, E. B.; Lu, F.; Portz, B.; Fisher, M. J.; Medellin, B. P.; Laremore, T. N.; Zhang, Y. J.; Gilmour, D. S.; Showalter, S. A. Phosphorylation induces sequence-specific conformational switches in the RNA polymerase II C-terminal domain. Nat Commun 2017, 8, 15233. DOI: 10.1038/ncomms15233 From NLM Medline.

(54) Martin, E. W.; Holehouse, A. S.; Grace, C. R.; Hughes, A.; Pappu, R. V.; Mittag, T. Sequence Determinants of the Conformational Properties of an Intrinsically Disordered Protein Prior to and upon Multisite Phosphorylation. J Am Chem Soc 2016, 138 (47), 15323-15335. DOI: 10.1021/jacs.6b10272 From NLM Medline.

(55) Qiao, Q.; Bowman, G. R.; Huang, X. Dynamics of an intrinsically disordered protein reveal metastable conformations that potentially seed aggregation. J Am Chem Soc 2013, 135 (43), 16092-16101. DOI: 10.1021/ja403147m From NLM Medline.

(56) Choi, U. B.; Sanabria, H.; Smirnova, T.; Bowen, M. E.; Weninger, K. R. Spontaneous Switching among Conformational Ensembles in Intrinsically Disordered Proteins. Biomolecules 2019, 9 (3). DOI: 10.3390/biom9030114 From NLM Medline.

(57) Samanta, H. S.; Chakraborty, D.; Thirumalai, D. Charge fluctuation effects on the shape of flexible polyampholytes with applications to intrinsically disordered proteins. J Chem Phys 2018, 149 (16), 163323. DOI: 10.1063/1.5035428 From NLM Medline.

(58) Schuler, B.; Borgia, A.; Borgia, M. B.; Heidarsson, P. O.; Holmstrom, E. D.; Nettels, D.; Sottini, A. Binding without folding – the biomolecular function of disordered polyelectrolyte complexes. Curr Opin Struc Biol 2020, 60, 66-76. DOI: 10.1016/j.sbi.2019.12.006 PMID - 31874413.

(59) Wang, W. Recent advances in atomic molecular dynamics simulation of intrinsically disordered proteins. Phys Chem Chem Phys 2021, 23 (2), 777-784. DOI: 10.1039/d0cp05818a From NLM Medline.

(60) Luong, T. D. N.; Nagpal, S.; Sadqi, M.; Munoz, V. A modular approach to map out the conformational landscapes of unbound intrinsically disordered proteins. Proc Natl Acad Sci U S A 2022, 119 (23), e2113572119. DOI: 10.1073/pnas.2113572119 From NLM Medline.

(61) Uversky, V. N. Multitude of binding modes attainable by intrinsically disordered proteins: a portrait gallery of disorder-based complexes. Chem Soc Rev 2011, 40 (3), 1623-1634. DOI: 10.1039/c0cs00057d From NLM Medline.

(62) Arai, M.; Sugase, K.; Dyson, H. J.; Wright, P. E. Conformational propensities of intrinsically disordered proteins influence the mechanism of binding and folding. Proceedings of the National Academy of Sciences 2015, 112 (31), 9614-9619. DOI: doi:10.1073/pnas.1512799112. Chu, X.; Nagpal, S.; Muñoz, V. Molecular Simulations of Intrinsically Disordered Proteins and Their Binding Mechanisms. In Protein Folding: Methods and Protocols, Muñoz, V. Ed.; Springer US, 2022; pp 343-362.

(63) Lazar, T.; Martínez-Pérez, E.; Quaglia, F.; Hatos, A.; Chemes, Lucía B.; Iserte, J. A.; Méndez, N. A.; Garrone, N. A.; Saldaño, Tadeo E.; Marchetti, J.;, et al. PED in 2021: a major update of the protein ensemble database for intrinsically disordered proteins. Nucleic Acids Research 2020, 49 (D1), D404-D411. DOI: 10.1093/nar/gkaa1021 (acccessed 5/5/2023).

(64) Crehuet, R.; Buigues, P. J.; Salvatella, X.; Lindorff-Larsen, K. Bayesian-Maximum-Entropy Reweighting of IDP Ensembles Based on NMR Chemical Shifts. Entropy 2019, 21 (9), 898. DOI: 10.3390/e21090898.

(65) Jones, D. T.; Cozzetto, D. DISOPRED3: precise disordered region predictions with annotated protein-binding activity. Bioinformatics 2015, 31 (6), 857-863. DOI: 10.1093/bioinformatics/btu744 From NLM Medline.

(66) Schrodinger, LLC. The PyMOL Molecular Graphics System, Version 1.8. 2015.

(67) Bonneau, R.; Tsai, J.; Ruczinski, I.; Chivian, D.; Rohl, C.; Strauss, C. E.; Baker, D. Rosetta in CASP4: progress in ab initio protein structure prediction. Proteins 2001, *Suppl* 5, 119-126. DOI: 10.1002/prot.1170 From NLM Medline.

(68) Rohl, C. A.; Strauss, C. E.; Misura, K. M.; Baker, D. Protein structure prediction using Rosetta. Methods Enzymol 2004, 383, 66-93. DOI: 10.1016/S0076-6879(04)83004-0 From NLM PubMed-not-MEDLINE.

(69) Rozycki, B.; Kim, Y. C.; Hummer, G. SAXS Ensemble Refinement of ESCRT-III CHMP3 Conformational Transitions. Structure 2011, 19 (1), 109-116. DOI: 10.1016/j.str.2010.10.006.

(70) Satopaa, V.; Albrecht, J.; Irwin, D.; Raghavan, B. Finding a “Kneedle” in a Haystack: Detecting Knee Points in System Behavior. In 2011 31st International Conference on Distributed Computing Systems Workshops, 20-24 June 2011, 2011; pp 166–171. DOI: 10.1109/ICDCSW.2011.20.

(71) Tesei, G.; Martins, J. M.; Kunze, M. B. A.; Wang, Y.; Crehuet, R.; Lindorff-Larsen, K. DEER-PREdict: Software for efficient calculation of spin-labeling EPR and NMR data from conformational ensembles. PLoS Comput Biol 2021, 17 (1), e1008551. DOI: 10.1371/journal.pcbi.1008551 From NLM Medline.

(72) Gillespie, J. R.; Shortle, D. Characterization of long-range structure in the denatured state of staphylococcal nuclease. I. Paramagnetic relaxation enhancement by nitroxide spin labels. J Mol Biol 1997, 268 (1), 158-169. DOI: 10.1006/jmbi.1997.0954 From NLM Medline.

(73) Ganguly, D.; Chen, J. Structural interpretation of paramagnetic relaxation enhancement-derived distances for disordered protein states. J Mol Biol 2009, 390 (3), 467-477. DOI: 10.1016/j.jmb.2009.05.019 From NLM Medline.

(74) Ward, J. H. Hierarchical Grouping to Optimize an Objective Function. J Am Stat Assoc 1963, 58 (301), 236-&. DOI: Doi 10.2307/2282967.

(75) Wishart, D. 256. Note: An Algorithm for Hierarchical Classifications. Biometrics 1969, 25 (1), 165- 170. DOI: 10.2307/2528688 (acccessed 2023/03/22/).JSTOR.

(76) Kalinin, S.; Peulen, T.; Sindbert, S.; Rothwell, P. J.; Berger, S.; Restle, T.; Goody, R. S.; Gohlke, H.; Seidel, C. A. A toolkit and benchmark study for FRET-restrained high-precision structural modeling. Nat Methods 2012, 9 (12), 1218-1225. DOI: 10.1038/nmeth.2222 From NLM Medline.

(77) Sindbert, S.; Kalinin, S.; Nguyen, H.; Kienzler, A.; Clima, L.; Bannwarth, W.; Appel, B.; Muller, S.; Seidel, C. A. Accurate distance determination of nucleic acids via Forster resonance energy transfer: implications of dye linker length and rigidity. J Am Chem Soc 2011, 133 (8), 2463-2480. DOI: 10.1021/ja105725e From NLM Medline.

(78) Dimura, M.; Peulen, T. O.; Hanke, C. A.; Prakash, A.; Gohlke, H.; Seidel, C. A. Quantitative FRET studies and integrative modeling unravel the structure and dynamics of biomolecular systems. Curr Opin Struct Biol 2016, 40, 163-185. DOI: 10.1016/j.sbi.2016.11.012 From NLM Medline.

(79) McGibbon, R. T.; Beauchamp, K. A.; Harrigan, M. P.; Klein, C.; Swails, J. M.; Hernandez, C. X.; Schwantes, C. R.; Wang, L. P.; Lane, T. J.; Pande, V. S. MDTraj: A Modern Open Library for the Analysis of Molecular Dynamics Trajectories. Biophys J 2015, 109 (8), 1528-1532. DOI: 10.1016/j.bpj.2015.08.015 From NLM Medline.

(80) Gebhardt, C.; Lehmann, M.; Reif, M. M.; Zacharias, M.; Gemmecker, G.; Cordes, T. Molecular and Spectroscopic Characterization of Green and Red Cyanine Fluorophores from the Alexa Fluor and AF Series. Chemphyschem 2021, 22 (15), 1546. DOI: 10.1002/cphc.202100509 From NLM PubMed-not- MEDLINE.

(81) Peulen, T. O.; Opanasyuk, O.; Seidel, C. A. M. Combining Graphical and Analytical Methods with Molecular Simulations To Analyze Time-Resolved FRET Measurements of Labeled Macromolecules Accurately. J Phys Chem B 2017, 121 (35), 8211-8241. DOI: 10.1021/acs.jpcb.7b03441 From NLM Medline.

(82) Grudinin, S.; Garkavenko, M.; Kazennov, A. Pepsi-SAXS: an adaptive method for rapid and accurate computation of small-angle X-ray scattering profiles. Acta Crystallographica Section D 2017, 73 (5), 449- 464. DOI: doi:10.1107/S2059798317005745.

(83) Neal, S.; Nip, A. M.; Zhang, H.; Wishart, D. S. Rapid and accurate calculation of protein 1H, 13C and 15N chemical shifts. Journal of Biomolecular NMR 2003, 26 (3), 215–240. DOI: 10.1023/A:1023812930288.

(84) Kay, L. E.; Keifer, P.; Saarinen, T. Pure absorption gradient enhanced heteronuclear single quantum correlation spectroscopy with improved sensitivity. Journal of the American Chemical Society 1992, 114 (26), 10663-10665. DOI: 10.1021/ja00052a088.

(85) Kay, L. E.; Torchia, D. A.; Bax, A. Backbone dynamics of proteins as studied by 15N inverse detected heteronuclear NMR spectroscopy: application to staphylococcal nuclease. Biochemistry 1989, 28 (23), 8972-8979. DOI: 10.1021/bi00449a003 From NLM.

