## Supplemental infromation for "Delineating Structural Propensities of the 4E-BP2 Protein via Integrative Modelling and Clustering"

#### 1. Conformer Generation

##### 1.1 FastFloppyTail (FFT)

FastFloppyTail is a PyRosetta-based algorithm that allows de novo predictions of protein structures across the spectrum of protein order including IDPs that contain folded domains [1]. Essentially, the method samples dihedral angles from a library of 3-mer residue fragments across the entire Protein Databank (PDB) and applies Rosetta energy minimization functions to build conformers for a given protein sequence. The 3-mer fragment library was generated using the Robetta web server [2]. The FFT algorithm requires that a certain fraction of disorder is assigned to each residue in the protein sequence. The fraction can be set by the user if regions within the sequence are well-known to be ordered or disordered (ex., from experimental data). In the absence of such information, disorder predictors that estimate per-residue disordered fractions based off the protein's amino acid sequence are used.

##### 1.2 Coil ensemble generation for 5P state

To concatenate N-terminal and C-terminal disordered region conformers generated from TraDES using the "coil" secondary structure option, the IDPConformerGenerator [3] library was used. Steric clash-free stitching was performed using functions within the recently developed local disordered region sampling (LDRS) module. To fulfil the requirements for the alignment and concatenation function, the disordered termini tails were generated with two additional overlapping residues to the folded domain. All 20 poses of the NMR structure 2MX4 of the folded domain were used equally throughout this process. A library of 20,000 TraDES coil structures each were used for the N-terminal IDR as well as the C-terminal IDR concatenation. To facilitate a maximum success rate during the steric-clash check process, sidechains from the TraDES coil structures were stripped prior to concatenation and later re-packed with Monte Carlo side-chain entropy (MC-SCE). After the automated alignment, clash-checking, concatenation, and sidechain re-packing, 1000 all-atom structures generated from each 20 NMR structure poses were used to form the final set of 20,000 full length 5P folded 4E-BP2 structures for analysis as the TraDES coil ensemble.

#### 2. Ensemble calculations

##### 2.1 Bayesian Maximum Entropy (BME)

The reduced chi-squared metric ($\chi^2_{red}$) quantifies the agreement between experimental data and back-calculated data $y_i^{EXP}$ and $y_i^{BC}(x_j)$ respectively, for the $i^{th}$ data point of the $j^{th}$ conformation:

$$\chi^2(\omega_1, \dots \omega_M) = \frac{1}{m}\sum_{i=1}^{m} \frac{\left(\sum_{j=1}^{M} \omega_j y_i^{BC}(x_j) - y_i^{EXP}\right)^2}{\sigma_i^2} \qquad (S1)$$

where $m$ is the number of experimental datapoints, $M$ is the number of conformations, $\omega_j$ is the optimized BME weight for conformation $j$ and $\sigma_i$ is the combined uncertainty from both experiment and back-calculation [4].

The relative Shannon entropy ($S_{REL}$) is a measure of the degree of BME optimization and is defined as:

$$S_{REL} = -\sum_{j=1}^{M} \omega_j \log \frac{\omega_j}{\omega_j^0} \qquad (S2)$$

where $\omega_j^0$ is the initial weight of conformation $j$ before BME optimization (i.e., $1/M$) [4].

A more intuitive metric of the degree of ensemble optimization is (in a qualitative sense) the "effective fraction of conformations", termed $N_{eff}$ [4].

$$N_{eff} = \exp(S_{REL}) \qquad (S3)$$

##### 2.2 BME applied to the 5P state

Experimental data measured between residues (CS, PRE) had all experimental data points removed that had both residues located inside the folded domain (residues 18-62). When these data were included, the BME refinement would not converge.

#### 2.3 Atomic Root-Mean-Square Deviation (RMSD)

The Root-Mean-Square Deviation (RMSD) between two protein structures $i$ and $j$ is computed using the following formula:

$$RMSD_{i,j} = \frac{1}{\sqrt{N}}\sqrt{\sum_{n=1}^{N}[\left(C_{x,n}^{(i)} - C_{x,n}^{(j)}\right)^2 + \left(C_{y,n}^{(i)} - C_{y,n}^{(j)}\right)^2 + \left(C_{z,n}^{(i)} - C_{z,n}^{(j)}\right)^2]} \qquad (S4)$$

where $N$ is the number of residues in a protein (121 for 4E-BP2) and $C_{w,n}^{(m)}$ is the $w$-coordinate position of the Cα atom on the $n^{th}$residue for conformer $m$. The algorithm samples various superimpositions of conformers $i$ and $j$ through a quaternion-based characteristic polynomial approach [5] and outputs the minimum RMSD of those computed.

#### 2.4 Hierarchical clustering linkage

Hierarchical clustering is performed using a pairwise inter-residue distance metric and a ward-variance minimization linkage through the SciPy Python package [6] [7]. This more sophisticated choice of linkage measure was chosen based on its robustness to noise in comparison to other commonly employed linkage methods [8]. The ward-variance linkage joins clusters by minimizing the increase in the error sum of squares [8]. This measure is defined as the following between two clusters $X$ and $Y$ for which when combined form the cluster $XY$:

$$R_{X,Y} = \sqrt{ESS_{XY} - (ESS_X + ESS_Y)} \qquad (S5)$$

where:

$$ESS_Z = \sum_{i=1}^{M_Z}\sum_{a=1}^{N(N-1)/2}\left|r_a^{(i)} - \bar{r}_a^{(Z)}\right|^2 \qquad (S6)$$

and $r_a^{(i)}$ is the distance between $C_\alpha$ atoms of the $a^{th}$ residue pair for the $i^{th}$ conformer in cluster $Z$, $M_Z$ is the number of conformers in cluster $Z$ and $N$ $is$ the number of residues. The column vector $\bar{r}_a^{(Z)}$ is the average distance between $C_\alpha$ atoms of the $a^{th}$ residue pair in cluster $Z$, and can be written as follows:

$$\bar{r}^{(Z)} = \begin{pmatrix} \sum_{i=1}^{M_Z} \frac{r_1^{(i)}}{M_Z} \\ \sum_{i=1}^{M_Z} \frac{r_2^{(i)}}{M_Z} \\ \vdots \\ \sum_{i=1}^{M_Z} \frac{r_{N(N-1)/2}^{(i)}}{M_Z} \end{pmatrix} \qquad (S7)$$

#### 2.5 Transforming dendrogram distance metric to Euclidean distance

To compare to other biophysical data, we convert the increase in the cluster error sum of squares (eq. S6) of combining clusters $X$ and $Y$ to form cluster $XY$ to the Euclidean distance between cluster means ($D_T$), defined as:

$$D_T = |\bar{r}^{(X)} - \bar{r}^{(Y)}| = \sqrt{\frac{M_{XY}}{M_X M_Y}} R_{X,Y} \qquad (S8)$$

where $|\bar{r}^{(X)} - \bar{r}^{(Y)}|$ is the Euclidean distance between the vectors of all non-degenerate (7260) mean inter-residue distance pairs for clusters $X$ ($\bar{r}^{(X)}$) and $Y$ ($\bar{r}^{(Y)}$). The variables $M_X, M_Y$ and $M_{XY}$ are the number of conformers in clusters $X$, $Y$ and $XY$, respectively. The normalized cutoff distance displayed in Figures S5A and S6A is this transformed distance divided by $\sqrt{7260}$ to yield a root-mean-squared deviation between pairwise inter-residue distances (rather than atomic coordinates).

#### 2.6 Global size and shape parameters

The weighted averages for $R_g$ and $A$ were back-calculated via the MDAnalysis Python package [9 10].

$$R_g = \sqrt{\frac{1}{2N^2}\sum_{i,j}^{N}\langle r_{ij}{}^2\rangle} \qquad (S9)$$

The $R_g$ is calculated as the root mean-squared sum over $C_\alpha$-$C_\alpha$ distances ($r_{ij}$), where $N$ is the number of $C_\alpha$ atoms (eq. S10). $A$ is calculated using the eigenvalues of the radius of gyration tensor (eq. S11).

$$A = 1 - \frac{3(\lambda_1\lambda_2 + \lambda_1\lambda_3 + \lambda_2\lambda_3)}{(\lambda_1 + \lambda_2 + \lambda_3)^2} \qquad (S10)$$

The $R_h$ was computed using both the Kirkwood-Riseman approximation [11] (eq. S12) and HYDROPRO [12] [13] with default parameters.

$$R_H{}^{-1} = \frac{1}{N^2}\sum_{i\neq j}\langle r_{ij}{}^{-1}\rangle \qquad (S11)$$

#### 2.7 Internal Scaling Profile (ISP)

The internal scaling profile $C_\alpha$ distance ($R_{|i-j|}$) is computed as the average distance between all residues that have a residue separation of $|i-j|$ and then as a weighted average over all conformations using BME weights for the optimized ensemble or $1/M$ weights for the TraDES coil ensemble. The fit to the internal scaling profile using eq. 5 was fitted in MATLAB by first transforming to a linear equation of the form:

$$\ln(R_{|i-j|}) = \nu \ln(|i-j|) + C \qquad (S12)$$

where $C = \ln(\sqrt{2l_p b})$ was fixed at $\ln(5.51)$ [14].

#### 2.8 RMSD of the Paramagnetic Relaxation Enhancement data

This PRE RMSD metric is formulated as follows:

$$PRE\ RMSD = \sqrt{\frac{\Sigma_{l=1}^{L} RMSD_l^2}{L}} \qquad (S13)$$

where:

$$RMSD_l = \sqrt{\frac{\sum_{k=1}^{K}\left(\hat{R}_l - R_l^{(k)}\right)^2}{K}} \qquad (S14)$$

where $K$ is the number of PRE data points for the labelling site located on the $l^{th}$ residue, $L$ is the number of PRE labelling sites, $\hat{R}_l$ and $R_l^{(i)}$ is the mean experimental PRE intensity ratio for the labelling site on the $l^{th}$ residue and the back-calculated value of the same labelling site for the $i^{th}$ conformer, respectively.

#### 2.9 Two-dimensional distance/contact maps

Two-dimensional mean inter-residue distance scaling maps were constructed by taking the weight averages ($\langle s_{ij}\rangle$) of the inter-residue $C_\alpha$ atom distances for each unique residue separation. The distances ($\langle s_{ij}\rangle_1$) were divided by the corresponding reference distances ($\langle s_{ij}\rangle_2$) at each unique residue separation to obtain the final scaling maps ($r_{i,j}^{norm} = \langle s_{ij}\rangle_1/\langle s_{ij}\rangle_2$).

$$\langle s_{ij}\rangle = \frac{\sum_{a=1}^{N(N-1)/2} w_a r_{ij,a}}{\sum_{a=1}^{N(N-1)/2} w_a} \qquad (S15)$$

Difference inter-residue contact maps were constructed by first taking the weighted fraction of conformers at each unique inter-residue separation with a $C_\alpha$-$C_\alpha$ cutoff distance of $< 8$ Å ($\langle f_{ij}\rangle$).

$$f_{ij} = \frac{\sum_{a=1}^{N(N-1)/2} w_a \, I_{\{r|r<8\,\text{Å}\}}(r_{ij,a})}{\sum_{a=1}^{N(N-1)/2} w_a} \qquad (S16)$$

where $I_A(a)$ is the indicator function for the set $A$ which outputs 1 if $a \in A$ and 0 otherwise. The fractions ($\langle f_{ij} \rangle_1$) were subtracted by the corresponding reference fractions ($\langle f_{ij} \rangle_2$) at each unique residue separation to obtain the final contact maps.

#### SUPPLEMENTAL FIGURES

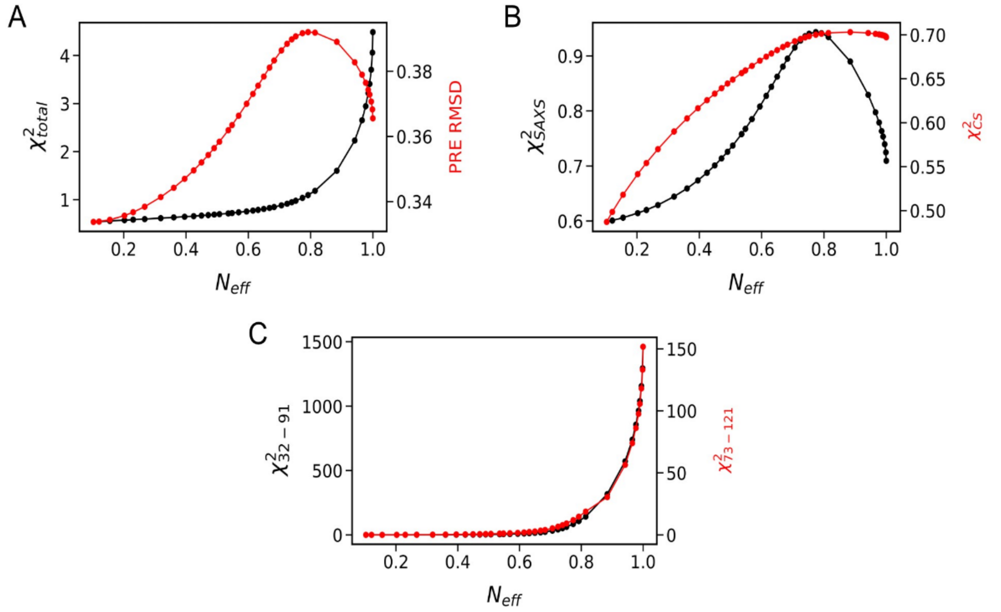


**Figure S1.** Restraint optimization of the NP 4E-BP2 ensemble including: (**A**) all restraints (smFRET, SAXS, CS) (black) and PRE RMSD (red), (**B**) SAXS (black) and CS (red), (**C**) 32-91 (black) and 73-121 (red) smFRET.

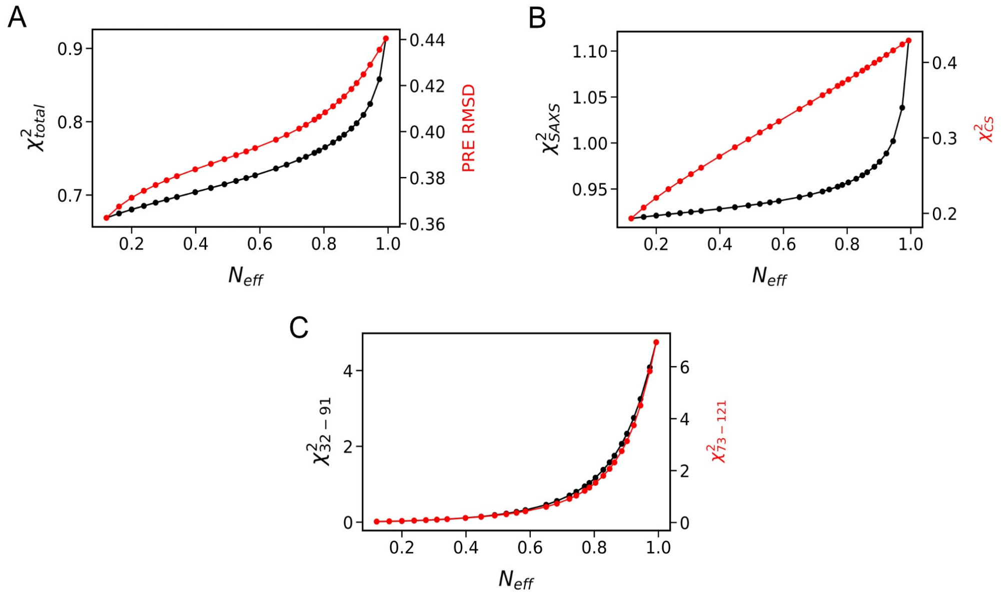


**Figure S2.** Restraint optimization of the 5P 4E-BP2 ensemble including: (**A**) all restraints (smFRET, SAXS, CS) (black) and PRE RMSD (red), (**B**) SAXS (black) and CS (red), (**C**) 32-91 (black) and 73-121 (red) smFRET.

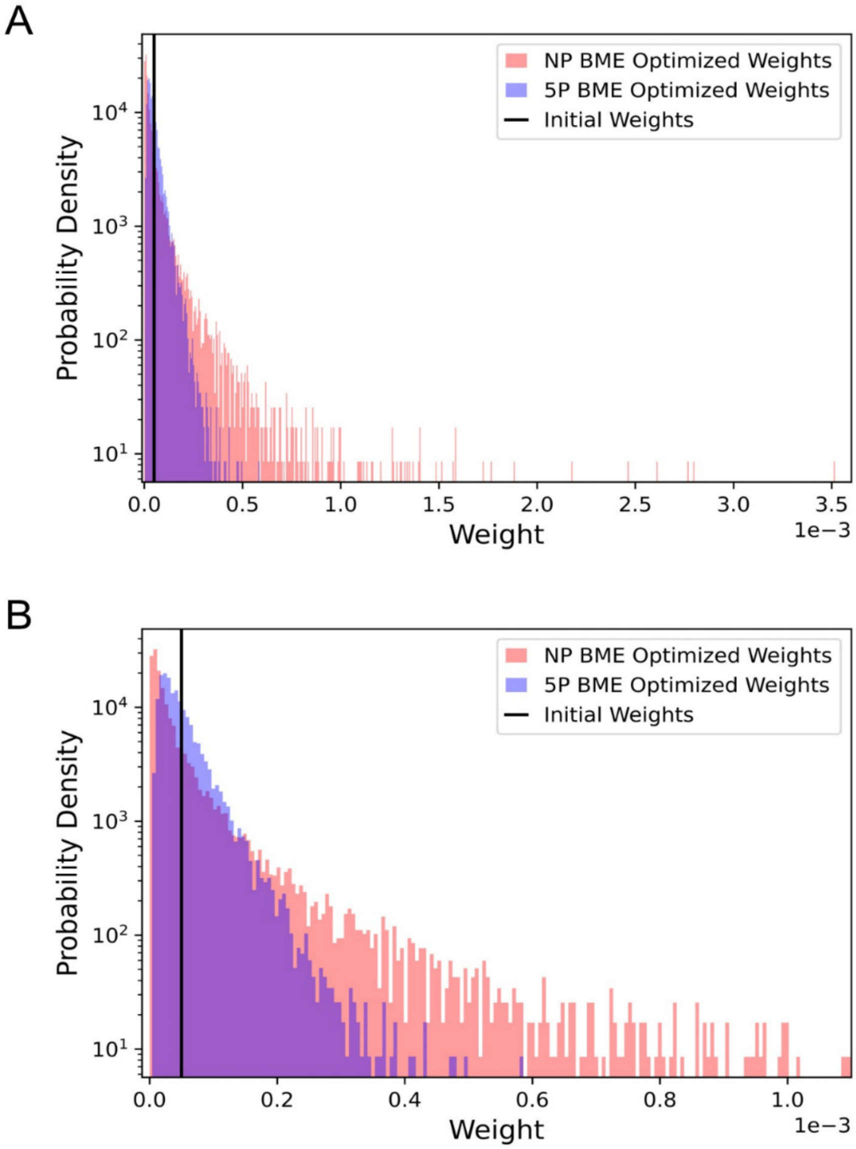


**Figure S3.** The distribution of conformer weights for the BME-optimized NP (red) and 5P (blue) 4E-BP2 ensembles. The unoptimized weight per conformer (1/20,000) is displayed as a black vertical line. Note that probability density is plotted on a log scale.

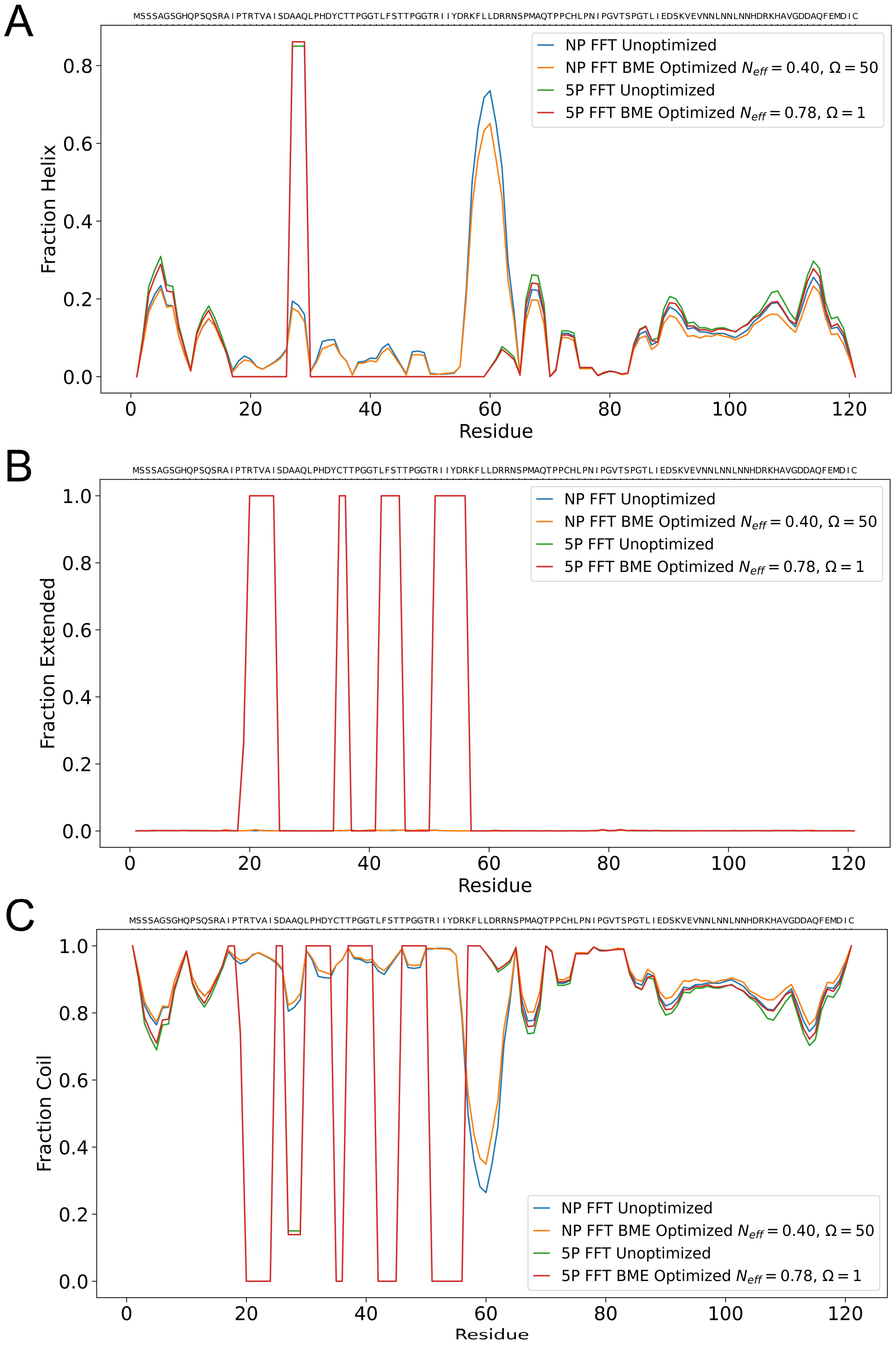


**Figure S4:** Secondary structure fractions of the BME-optimized NP and 5P 4E-BP2 ensembles calculated using the Dictionary of Protein Secondary Structure (DSSP) method [15] implemented in the MDTraj Python package [16]. Three secondary structure types are displayed: (**A**) helical (including alpha helix, 3-helix and 5 helix), (**B**) extended (including beta-bridge and beta ladder) and (**C**) coil.

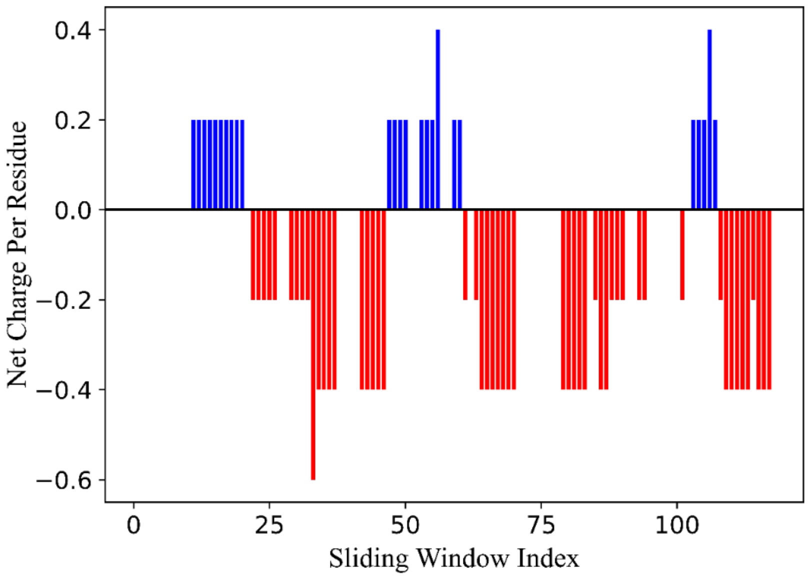


**Figure S5:** NCPR calculated using a five-residue sliding-window [17] across the sequence of 5P 4E-BP2. The charge of each amino acid was taken to be the most populous charge state at pH 7 using pKa of each amino acid [18], and phosphate groups were assigned a -2 charge.

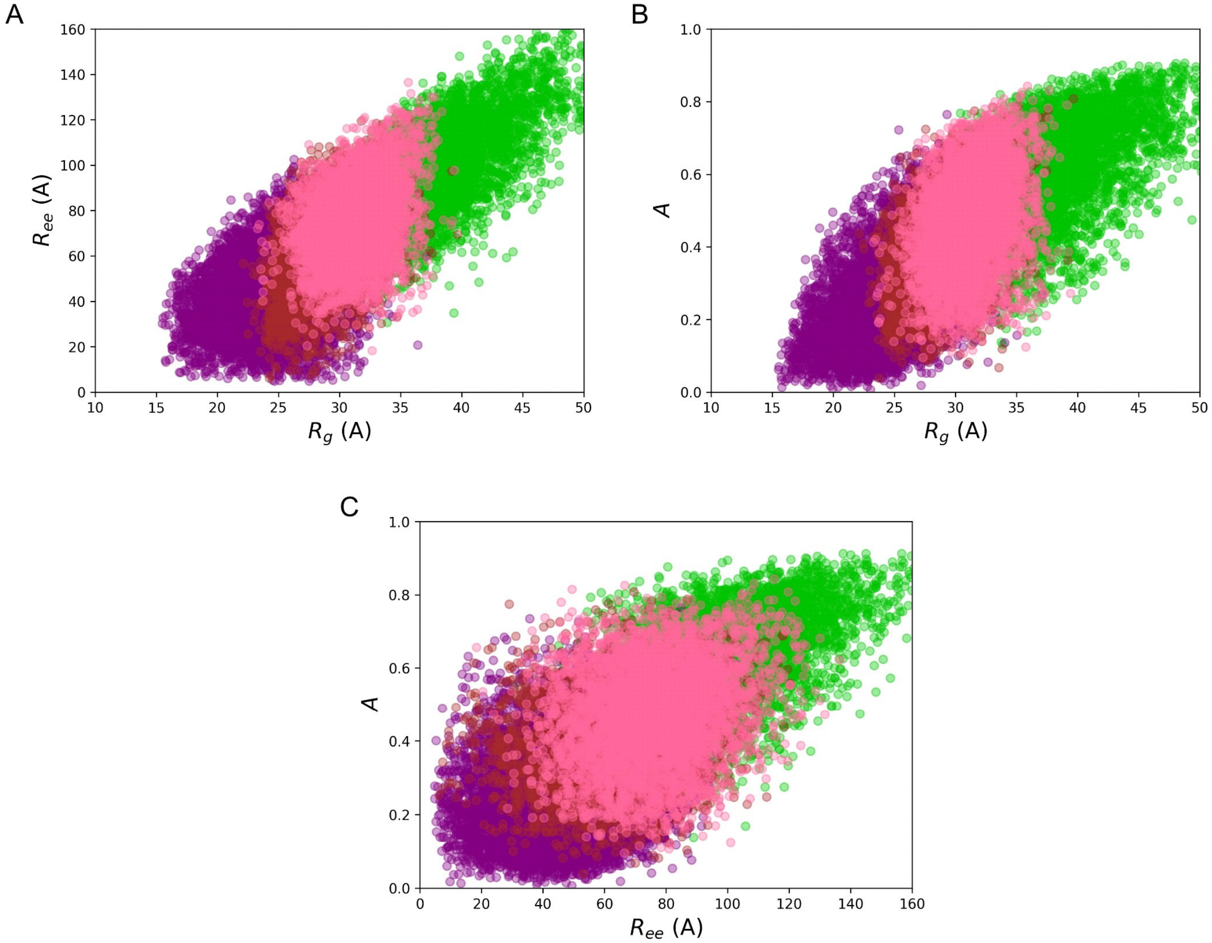


**Figure S6:** Global size and shape parameters of the unrestrained NP FFT ensemble. The data is colored according to the cluster assignments depicted in **Fig. 5A** in the main text: Cluster 1 (green), Cluster 2 (purple), Cluster 3 (brown) and Cluster 4 (pink). The relations plotted are the end-to-end distance *vs.* the radius of gyration (**A**), asphericity *vs.* the radius of gyration (**B**) and asphericity *vs.* the end-to-end distance (**C**).

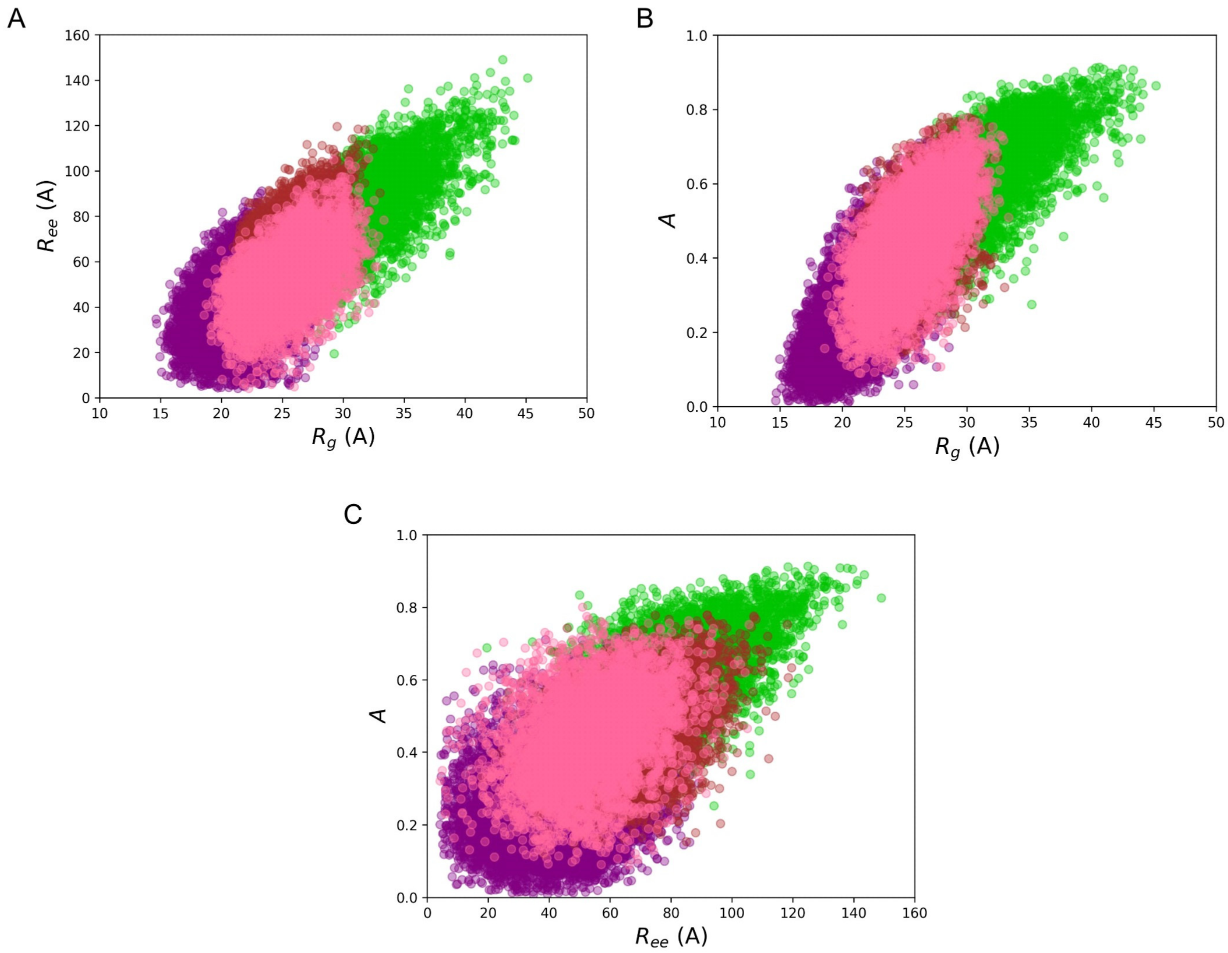


**Figure S7:** Global size and shape parameters of the unrestrained 5P FFT ensemble. The data is colored according to the cluster assignments depicted in **Fig. 6A** in the main text: Cluster 1 (green), Cluster 2 (purple), Cluster 3 (brown) and Cluster 4 (pink). The relations plotted are the end-to-end distance *vs.* the radius of gyration (**A**), asphericity *vs.* the radius of gyration (**B**) and asphericity *vs.* the end-to-end distance (**C**).

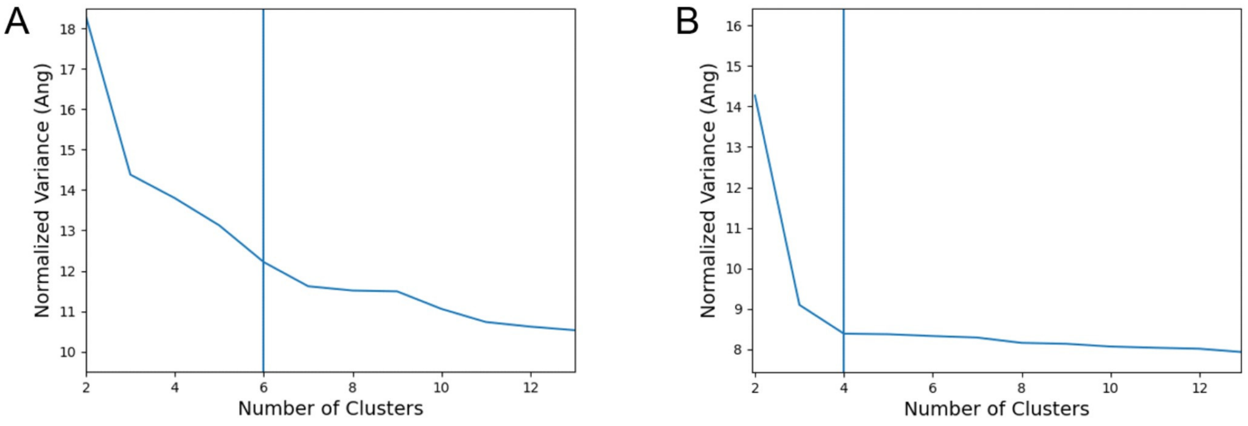


**Figure S8.** Normalized variance for determining the number of clusters in the BME-optimized NP **(A)** and 5P **(B)** 4E-BP2 conformational ensembles.

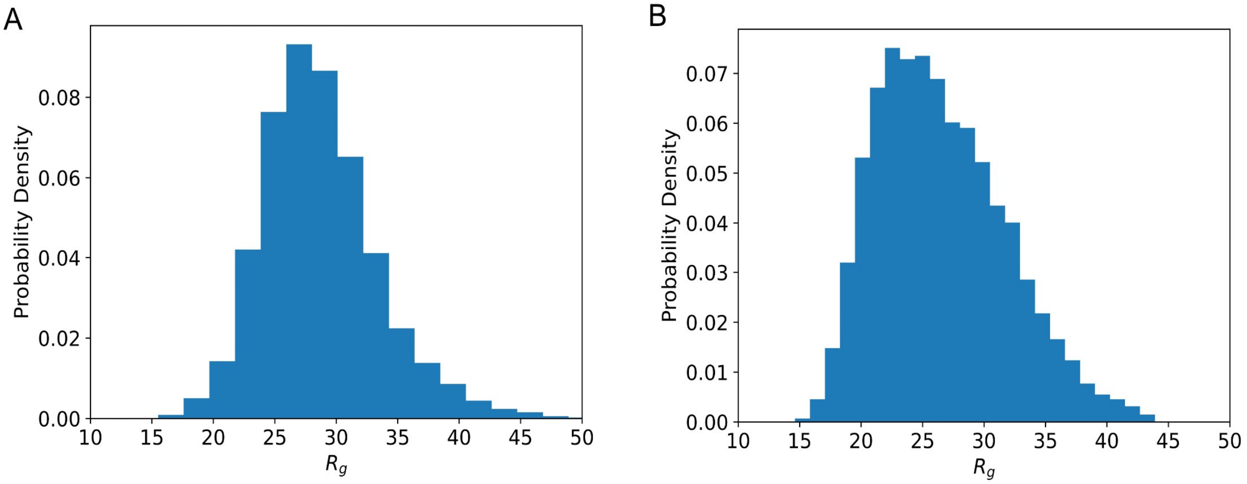


**Figure S9:** Probability distributions of the radius of gyration for the BME-optimized NP (**A**) and 5P (**B**) ensembles.

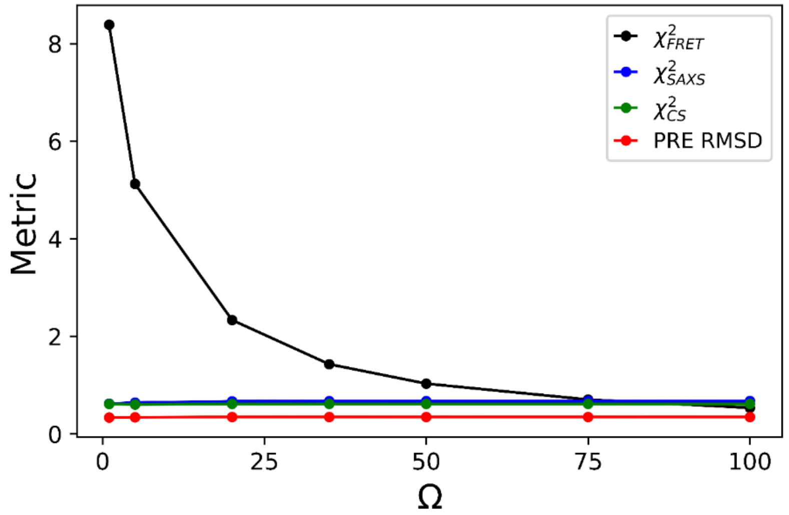


**Figure S10:** Varying the hyperparameter Ω (weight of the smFRET restraint) in BME optimization of the NP 4E-BP2 ensemble. PRE RMSD validation and reduced chi-squared values for smFRET, SAXS, and CS were back-calculated for Ω values from 1 to 100. The optimal $\theta$ for each optimization was chosen as the "knee" point of the PRE RMSD validation curve. The fitting to smFRET data improves significantly with negligible increases to other fits as Ω increases. At $\Omega = 50$ the FRET data is fit with $\chi^2_{FRET} = 1.03$, as such this value was selected in the final optimization.

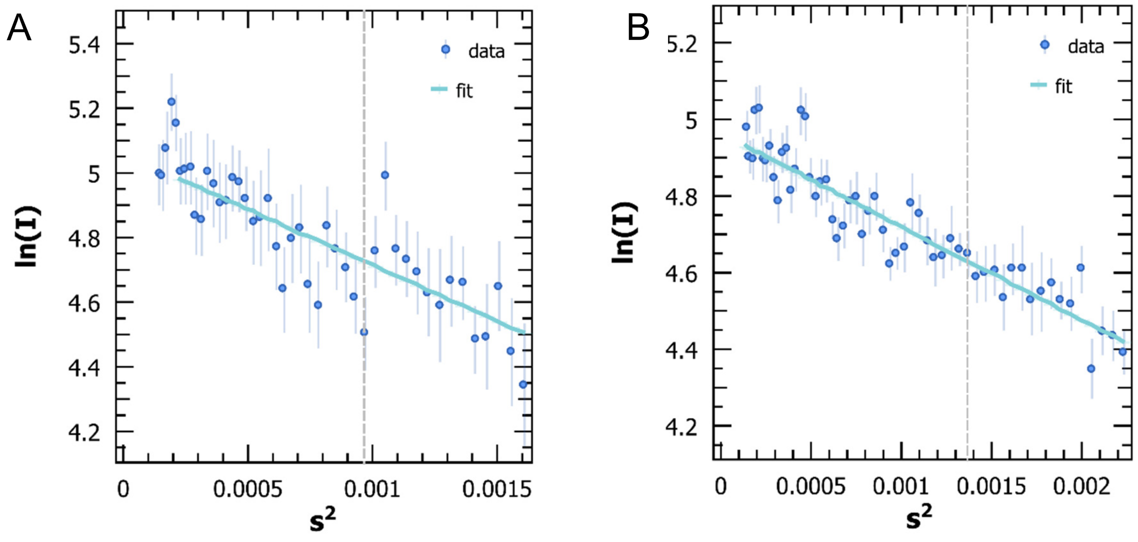


**Figure S11:** Guinier plots (data and fit) of NP 4E-BP2 (**A**) and 5P 4E-BP2 (**B**). The SAXS data was acquired using an AntonPaar instrument (see Methods), and fitted using the AutoRg function in the PRIMUS program [19]. The resulting radii of gyration are 32.18 ± 2.22 Å and 27.06 ± 0.69 Å for NP and 5P states, respectively. Experimental uncertainties for each data point are shown as vertical lines.

#### SUPPLEMENTAL TABLES

**Table S1.** Fitting metrics of the unoptimized and the BME-optimized NP 4E-BP2 ensembles.

| | **Unoptimized** | **BME Optimized** ($\Omega = 50,\ N_{eff} = 0.40, \theta = 35$) |
|---|---|---|
| $\chi^2_{total}$ | 4.5 | 0.64 |
| $\chi^2_{SAXS}$ | 0.71 | 0.67 |
| $\chi^2_{CS}$ | 0.70 | 0.62 |
| $\chi^2_{FRET}$ | 806 | 1.0 |
| $\chi^2_{32-91}$ | 1461 | 1.7 |
| $\chi^2_{73-121}$ | 152 | 0.39 |
| **PRE RMSD** | 0.37 | 0.35 |

**Table S2.** Fitting metrics of the unoptimized and the BME-optimized 5P 4E-BP2 ensembles.

| | **Unoptimized** | **BME Optimized** ($\Omega = 1,\ N_{eff} = 0.78, \theta = 27$) |
|---|---|---|
| $\chi^2_{total}$ | 0.99 | 0.76 |
| $\chi^2_{SAXS}$ | 1.22 | 0.95 |
| $\chi^2_{CS}$ | 0.43 | 0.37 |
| $\chi^2_{FRET}$ | 6.4 | 1.2 |
| $\chi^2_{32-91}$ | 5.1 | 1.0 |
| $\chi^2_{73-121}$ | 7.6 | 1.3 |
| **PRE RMSD** | 0.44 | 0.41 |

**Table S3.** Fitting parameters to the power-law ISP scaling for intermediate- and long-range residue separations in NP 4E-BP2 ensembles.*

| | $l_p$ **(Å)** | $b$ **(Å)** | $\nu$ |
|---|---|---|---|
| **NP FFT BME (θ = 35, Ω = 50) Short Scaling Fit** | 4 | 3.8 | 0.556 |
| **NP FFT BME (θ = 35, Ω = 50) Long Scaling Fit** | 4 | 3.8 | 0.539 |
| **NP TraDES RC Unrestrained Global Scaling Fit** | 4 | 3.8 | 0.561 |

* Short range is defined as $10 \le |i - j| \le 40$ and long range as $100 \le |i - j| \le 120$.

**Table S4.** NP 4E-BP2 sequence-charge parameters.*

| NCPR | FCR | Fraction of positive residues | Fraction of negative residues | Hydropathy | κ |
|---|---|---|---|---|---|
| -0.0167 | 0.1833 | 0.0833 | 0.1000 | 3.9117 | 0.1552 |

*Back-calculations of all parameters were computed using the CIDER program [20].

**Table S5.** Sum of conformer weights for NP 4E-BP2 clusters before and after applying BME-derived weights.

| Cluster # | Initial | Optimized |
|---|---|---|
| **1** | 0.225 | 0.115 |
| **2** | 0.409 | 0.428 |
| **3** | 0.162 | 0.266 |
| **4** | 0.204 | 0.191 |

**Table S6.** Sum of conformer weights for 5P 4E-BP2 clusters before and after applying BME-derived weights.

| Cluster # | Initial | Optimized |
|---|---|---|
| **1** | 0.201 | 0.277 |
| **2** | 0.413 | 0.331 |
| **3** | 0.179 | 0.161 |
| **4** | 0.207 | 0.231 |

**Table S7.** Optimization parameters and experimental back-calculated biophysical parameters of NP 4E-BP2 for four independent initial pools of 5000 conformers.*

| FFT Pool Number | $N_{eff}$ | PRE RMSD | $\chi^2_{total}$ | $\chi^2_{SAXS}$ | $\chi^2_{CS}$ | $\langle E \rangle_{32-91}$ | $\langle E \rangle_{73-121}$ |
|---|---|---|---|---|---|---|---|
| **1** | 0.39 | 50 | 0.35 | 0.64 | 0.67 | 0.61 | 0.62 |
| **2** | 0.40 | 50 | 0.33 | 0.65 | 0.68 | 0.63 | 0.62 |
| **3** | 0.43 | 50 | 0.34 | 0.66 | 0.69 | 0.63 | 0.62 |
| **4** | 0.39 | 50 | 0.35 | 0.65 | 0.67 | 0.62 | 0.62 |

*Optimization parameters ($N_{eff}$ and Ω), external validation (PRE RMSD), and agreement with restraint data ($\chi^2_{total}$, $\chi^2_{SAXS}$, $\chi^2_{CS}$, $\langle E \rangle_{32-91}$ and $\langle E \rangle_{73-121}$) for the BME refinement of 4 independently calculated NP 4E-BP2 FFT ensembles. Uncertainties of back-calculated FRET efficiencies are ±0.03 (see Methods 4.4)

**Table S8.** Optimization parameters and experimental back-calculated biophysical parameters of 5P 4E-BP2 for four independent initial pools of 5000 conformers.*

| FFT Pool Number | $N_{eff}$ | PRE RMSD | $\chi^2_{total}$ | $\chi^2_{SAXS}$ | $\chi^2_{CS}$ | $\langle E \rangle_{32-91}$ | $\langle E \rangle_{73-121}$ |
|---|---|---|---|---|---|---|---|
| **1** | 0.78 | 0.39 | 0.76 | 0.95 | 0.38 | 0.48 | 0.48 |
| **2** | 0.78 | 0.39 | 0.76 | 0.96 | 0.37 | 0.48 | 0.48 |
| **3** | 0.83 | 0.39 | 0.77 | 0.96 | 0.38 | 0.48 | 0.48 |
| **4** | 0.81 | 0.40 | 0.76 | 0.96 | 0.38 | 0.48 | 0.48 |

*Optimization parameters ($N_{eff}$ and Ω), external validation (PRE RMSD), and agreement with restraint data ($\chi^2_{total}$, $\chi^2_{SAXS}$, $\chi^2_{CS}$, $\langle E \rangle_{32-91}$ and $\langle E \rangle_{73-121}$) for the BME refinement of 4 independently calculated 5P 4E-BP2 FFT ensembles (with $\Omega = 1$). Uncertainties of back-calculated FRET efficiencies are ±0.03 (see Methods 4.4).

### References


(1) Ferrie, J. J.; Petersson, E. J. A unified de novo approach for predicting the structures of ordered and disordered proteins. *The Journal of Physical Chemistry B* **2020**, *124* (27), 5538-5548.
(2) Kim, D. E.; Chivian, D.; Baker, D. Protein structure prediction and analysis using the Robetta server. *Nucleic Acids Res* **2004**, *32* (Web Server issue), W526-531. DOI: 10.1093/nar/gkh468 From NLM Medline.
(3) Teixeira, J. M.; Liu, Z. H.; Namini, A.; Li, J.; Vernon, R. M.; Krzeminski, M.; Shamandy, A. A.; Zhang, O.; Haghighatlari, M.; Yu, L. IDPConformerGenerator: A Flexible Software Suite for Sampling the Conformational Space of Disordered Protein States. *The Journal of Physical Chemistry A* **2022**, *126* (35), 5985-6003.
(4) Bottaro, S.; Bengtsen, T.; Lindorff-Larsen, K. Integrating Molecular Simulation and Experimental Data: A Bayesian/Maximum Entropy Reweighting Approach. *Methods Mol Biol* **2020**, *2112*, 219-240. DOI: 10.1007/978-1-0716-0270-6_15 From NLM Medline.
(5) Theobald, D. Rapid calculation of RMSDs using a quaternion-based characteristic polynomial. *Acta Crystallographica Section A* **2005**, *61* (4), 478-480. DOI: doi:10.1107/S0108767305015266.
(6) Bar-Joseph, Z.; Gifford, D. K.; Jaakkola, T. S. Fast optimal leaf ordering for hierarchical clustering. *Bioinformatics* **2001**, *17 Suppl 1*, S22-29. DOI: 10.1093/bioinformatics/17.suppl_1.s22 From NLM Medline.
(7) Virtanen, P.; Gommers, R.; Oliphant, T. E.; Haberland, M.; Reddy, T.; Cournapeau, D.; Burovski, E.; Peterson, P.; Weckesser, W.; Bright, J.; et al. SciPy 1.0: fundamental algorithms for scientific computing in Python. *Nat Methods* **2020**, *17* (3), 261-272. DOI: 10.1038/s41592-019-0686-2 From NLM Medline.
(8) Ward, J. H. Hierarchical Grouping to Optimize an Objective Function. *J Am Stat Assoc* **1963**, *58* (301), 236-&. DOI: Doi 10.2307/2282967.
(9) Michaud-Agrawal, N.; Denning, E. J.; Woolf, T. B.; Beckstein, O. MDAnalysis: a toolkit for the analysis of molecular dynamics simulations. *J Comput Chem* **2011**, *32* (10), 2319-2327. DOI: 10.1002/jcc.21787 From NLM Medline.
(10) Gowers, R.; Linke, M.; Barnoud, J.; Reddy, T.; Melo, M.; Seyler, S.; Domański, J.; Dotson, D.; Buchoux, S.; Kenney, I.; et al. MDAnalysis: A Python Package for the Rapid Analysis of Molecular Dynamics Simulations. 2016, 2016; SciPy. DOI: 10.25080/majora-629e541a-00e.
(11) Kirkwood, J. G.; Riseman, J. The intrinsic viscosities and diffusion constants of flexible macromolecules in solution. *The Journal of Chemical Physics* **1948**, *16* (6), 565-573.
(12) García De La Torre, J.; Huertas, M. L.; Carrasco, B. Calculation of Hydrodynamic Properties of Globular Proteins from Their Atomic-Level Structure. *Biophysical Journal* **2000**, *78* (2), 719-730. DOI: 10.1016/s0006-3495(00)76630-6.
(13) Ortega, A.; Amoros, D.; Garcia de la Torre, J. Prediction of hydrodynamic and other solution properties of rigid proteins from atomic- and residue-level models. *Biophys J* **2011**, *101* (4), 892-898. DOI: 10.1016/j.bpj.2011.06.046 From NLM Medline.
(14) Gomes, G. N. W.; Krzeminski, M.; Namini, A.; Martin, E. W.; Mittag, T.; Head-Gordon, T.; Forman-Kay, J. D.; Gradinaru, C. C. Conformational Ensembles of an Intrinsically Disordered Protein Consistent with NMR, SAXS, and Single-Molecule FRET. *Journal of the American Chemical Society* **2020**, *142* (37), 15697-15710. DOI: 10.1021/jacs.0c02088.
(15) Kabsch, W.; Sander, C. Dictionary of protein secondary structure: pattern recognition of hydrogen-bonded and geometrical features. *Biopolymers* **1983**, *22* (12), 2577-2637. DOI: 10.1002/bip.360221211 From NLM Medline.
(16) McGibbon, R. T.; Beauchamp, K. A.; Harrigan, M. P.; Klein, C.; Swails, J. M.; Hernandez, C. X.; Schwantes, C. R.; Wang, L. P.; Lane, T. J.; Pande, V. S. MDTraj: A Modern Open Library for the Analysis of Molecular Dynamics Trajectories. *Biophys J* **2015**, *109* (8), 1528-1532. DOI: 10.1016/j.bpj.2015.08.015 From NLM Medline.

(17) Holehouse, A. S.; Das, R. K.; Ahad, J. N.; Richardson, M. O.; Pappu, R. V. CIDER: Resources to Analyze Sequence-Ensemble Relationships of Intrinsically Disordered Proteins. *Biophys J* **2017**, *112* (1), 16-21. DOI: 10.1016/j.bpj.2016.11.3200  From NLM Medline.
(18) Haynes, W. *CRC Handbook of Chemistry and Physics, 92nd Edition*; 2011. DOI: 10.1201/b17379.
(19) Manalastas-Cantos, K.; Konarev, P. V.; Hajizadeh, N. R.; Kikhney, A. G.; Petoukhov, M. V.; Molodenskiy, D. S.; Panjkovich, A.; Mertens, H. D. T.; Gruzinov, A.; Borges, C.; et al. ATSAS 3.0: expanded functionality and new tools for small-angle scattering data analysis. *Journal of Applied Crystallography* **2021**, *54* (1), 343-355. DOI: doi:10.1107/S1600576720013412.
(20) Das, R. K.; Pappu, R. V. Conformations of intrinsically disordered proteins are influenced by linear sequence distributions of oppositely charged residues. *Proceedings of the National Academy of Sciences* **2013**, *110* (33), 13392-13397. Holehouse, A. S.; Das, R. K.; Ahad, J. N.; Richardson, M. O.; Pappu, R. V. CIDER: resources to analyze sequence-ensemble relationships of intrinsically disordered proteins. *Biophysical journal* **2017**, *112* (1), 16-21.
